# Environmental DNA analysis needs local reference data to inform taxonomy-based conservation policy – A case study from Aotearoa / New Zealand

**DOI:** 10.1101/2021.10.22.465527

**Authors:** Paul Czechowski, Michel de Lange, Michael Heldsinger, Will Rayment, Christopher Hepburn, Monique Ladds, Michael Knapp

## Abstract

Effective management of biodiversity requires regular surveillance of multiple species. Analysis of environmental DNA by metabarcoding (eDNA) holds promise to achieve this relatively easily. However, taxonomic inquiries into eDNA data need suitable molecular reference data, which are often lacking. We evaluate the impact of this reference data void in a case study of fish diversity in the remote fiords of New Zealand. We compared eDNA-derived species identifications against Baited Remote Underwater Video (BRUV) data collected at the same time and locations as the eDNA data. Furthermore, we cross referenced both eDNA and BRUV data against species lists for the same region obtained from literature surveys and the Ocean Biodiversity Information System (OBIS). From all four data sources, we obtained a total of 116 species records (106 ray-finned fishes, 10 cartilaginous fishes; 59 from literature, 44 from eDNA, 25 from BRUV, 25 from OBIS). Concordance of taxonomies between the data sources dissolved with lowering taxonomic levels, most decisively so for eDNA data. BRUV agreed with local biodiversity information much better and fared better in detecting regional biodiversity dissimilarities. We provide evidence that eDNA metabarcoding will remain a powerful but impaired tool for species-level biodiversity management without locally generated reference data.

## Introduction

Marine reserve (MR) networks conserve biodiversity by stabilizing communities and maintaining food web structure (Wing & Jack, 2013). Effective management of MR biodiversity requires regular surveillance, for example to avoid overexploitation by fishing (Wing & Jack, 2013), or to avoid damage through influx of non-indigenous species (Cunningham, 2019). Fish surveillance is of particular interest due to their sensitivity to most forms of human disturbance, their usefulness at all levels of biological organization, and the favourable benefit-to-cost ratio of fish assessment programmes (Harris, 1995).

Analysis of environmental DNA (eDNA) metabarcoding data is a well-established molecular technique for multispecies surveys (Cristescu & Hebert, 2018). Environmental DNA metabarcoding holds promise for biodiversity surveys intended to inform biodiversity management – associated techniques are regarded more cost-efficient than traditional methods (such as baited remote underwater video surveys – BRUV), less dependent on expert taxonomic knowledge, can be standardized, and are able to inform on a broad range of taxa (Sigsgaard et al., 2020).

Reliable low-level taxonomic annotation is a prerequisite for useful biodiversity management and biological surveillance (e.g., Jack & Wing 2013). For example, in a southern New Zealand context, *Parapercis colias* (blue cod) is of high commercial interest, but three other of the 79 cod species are known from New Zealand (Roberts et al., 2019), so that genus information alone is ambiguous for determining blue cod presence or absence. Accordingly, higher-level taxonomic classifications (e.g., family, and order levels) are even less informative for species level conservation management. This reality translates into the desire for obtaining perfect 100 bp to 200 bp alignments (Huson et al. 2007) between an unknown eDNA-derived query sequence and a well described reference sequence derived from a valid species. In practice, absence of such reference data necessitates relaxation of taxonomy-assigning alignment parameters to retain sufficient eDNA data for analysis, and in consequence the data’s informative quality suffers.

Availability of suitable reference data for metabarcoding is highly variable depending on taxonomic groups and geographic locations, with fish considered relatively well covered in Barcode of Life Data Systems (BOLD) and NCBIs GenBank (Benson et al. 2011) for some regions such as Europe (Weigand et al., 2019). Arguably, fewer reference data are available for fish of southern New Zealand. For example, for six commonly used 12S primer pairs, recognized as well suitable for fish multispecies surveys (Weigand et al., 2019), an average of 36% of all northern European fish species are available as reference data, but only 26% of southern New Zealand species (GAPeDNA v1.0.1 web interface, 11-Sep-2021; Marques et al. 2021; also see SI Table 1).

In this study, we evaluate the impact of taxonomic data limitations on multispecies surveys using the example of fish in the UNESCO World Heritage Site Te Wahipounamu (Fiordland) in southern New Zealand. We compare the results of concurrent eDNA and BRUV surveys and cross-reference these data against species lists for the same region obtained from literature and the Ocean Biodiversity Information System (OBIS). The fish diversity of Te Wahipounamu has been described based on a diverse range of mostly visual methods. If our BRUV and eDNA approaches work optimally, we should see a strong overlap between these field data and previously described fish diversity records of the region.

## Methods

For this study, we evaluated the presence of Actinopterygii (ray-finned fishes) and Chondrichthyes (cartilaginous fishes) species in one MR, two commercial exclusion zones (all “MR”), and corresponding control areas in southern Te Wahipounamu, New Zealand (west coast, approximately from −44.3 to −46.25 Southern latitude; Fig. 1a). We obtained and analysed eDNA and BRUV data as well as electronic records proximate to the field work area from the Ocean Biodiversity Information System (OBIS; Ausubel 1999). Furthermore, a reference list of ray-finned fishes and cartilaginous fishes that have been observed in Fiordland was assembled from literature. All observations were formalized using NCBI taxonomy (Federhen, 2012), including trivial names, and limited to classes Actinopterygii and Chondrichthyes.

**Fig. 1:**
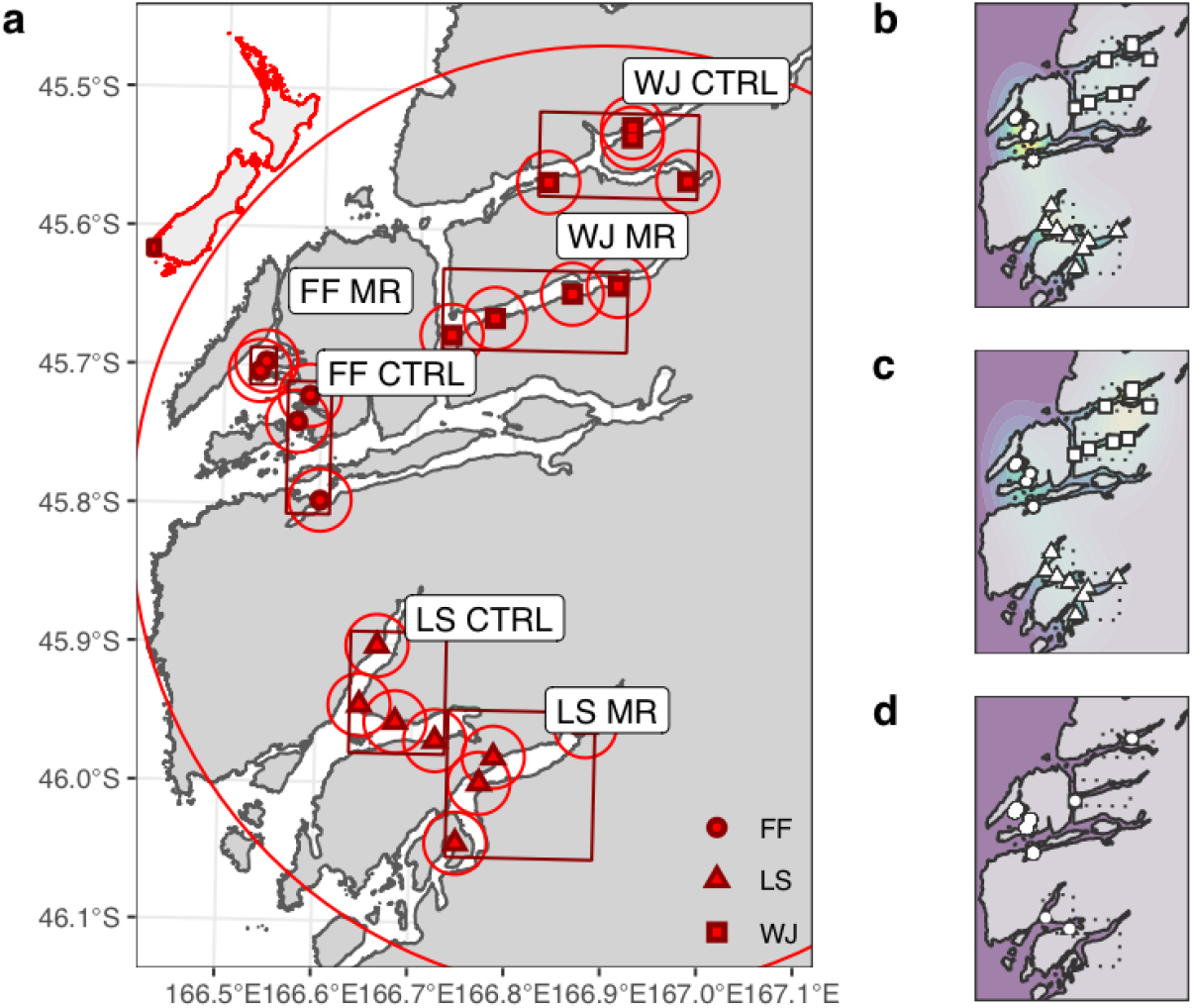
Field work area, description, sites, and data coverage for eDNA, BRUV, and OBIS data. **a:** We obtained biodiversity information from baited remote underwater video (BRUV) footage and environmental DNA (eDNA) data from 21 field work sites across three sampling regions (highlighted by rectangles) – Five Fingers (FF), Long Sound (LS) and Wet Jacket (WJ). In each region we collected samples inside marine reserves / commercial exclusion zones (MR) and outside in control areas (CTRL). To obtain additional biodiversity information, we queried the Ocean Biodiversity Information System (OBIS – https://obis.org/) for records within a 2.5 km radius of each field work site (small circles) for the purpose of community structure analysis. Furthermore, we obtained OBIS records for the entire sampling region (large circle) to extend our species list alongside species mentioned across various literature sources (Table 1, SI Table 3). **b:** Environmental DNA (eDNA), and **c:** BRUV data in a spatial context, lighter colour indicates a higher density of distinct species observations (corresponding to numerical values in Fig. 2). **d:** Species data for all filed work sites could not be obtained from OBIS, necessitating the exclusion of this data in the statistical analyses of regional biodiversity data. Graph created using R package *ggplot2 (3.3.5)*.

Literature data, itself obtained using various methods, were extracted from five sources, including one meta-analysis (SI Table 2). OBIS data were downloaded for a 38 km radius around all field work sites (centre point W 166.89°, S −45.80°), as well as for smaller areas surrounding individual field work sites, (2.5 km radius; Fig. 1a).

For a detailed description of field and laboratory work please refer to SI. For eDNA collection and BRUV filming we visited three locations in southern Te Wahipounamu (Moana Uta / Wet Jacket Arm, Taumoana / Five Fingers, and Te Tapuwae a Hua / Long Sound; henceforth WJ MR, FF MR, and LS MR), and accompanying control areas outside those MRs (henceforth WJ CTRL, FF CTRL, and LS CTRL), from 12.–22. December 2019 (Fig. 1a.) Within each sampling location, at randomised sites, we collected eDNA (mean depth 14.05 m, med.: 15, sd.: 1.4 m), and subsequently deployed BRUV assemblies (mean depth 15.6 m, med.: 16, sd.: 2.6 m; SI). We considered data from 21 sites (FF: 2 FF MR and 3 FF CTRL, WJ: 4 WJ MR and 4 WJ CTRL, and LS: 4 LS MR and 4 LS CTRL). We collected two 900 ml water samples with eDNA at each site, filtered them alongside negative controls, then sealed and stored them until further processing. BRUV footage was obtained for one hour and analysed by eye.

Environmental DNA was isolated in a PCR-free facility alongside extraction and cross-contamination controls (SI: four species of tropical freshwater fish). After *in silico* PCR (SI), we amplified our extracts with the well-established and widely used 12S MiFish primers (“MiFish-U”; Miya et al. 2015; see SI Table 1 for primer comparison), targeting Actinopterygii. Chondrichthyes were targeted with slightly altered derivatives (“Elas02”, Taberlet et al. 2018). Our single-step PCRs were cycled 45 times, with annealing temperatures of 45 °C (MiFish-U) or 40 °C (Elas02). Amplified eDNA was then pooled, visualised, purified, combined equimolarly, diluted to 4.5 pmol, and sequenced on an Illumina MiSeq (Illumina, San Diego, US-CA; kit v2, 300 cycles, single-ended).

We defined Amplicon Sequence Variants (ASVs; Callahan et al. 2017) from eDNA after demultiplexing with Cutadapt v3.0 (Martin, 2011), using Qiime2 2020-08 (Bolyen et al., 2019) and DADA2 1.10.0 (Callahan et al., 2016). To yield high quality sequence data we did not allow any mismatches, nor Expected Errors (Edgar & Flyvbjerg, 2015) during demultiplexing. Taxonomic annotation of denoised data was obtained using Blast 2.10.0+ (Camacho et al., 2009) and a local full copy of the NCBI nucleotide collection (April 2020; Benson et al., 2011) while excluding environmental samples. To yield a maximum of taxonomically annotated ASVs, we chose relaxed taxonomic assignment parameters in combination with an e-value to retain only the most significant alignments. We required a minimum identity of 75% among all alignments and kept five high-scoring pairs for each eDNA query, each of which needed a minimum coverage of 95% to be retained. The minimal acceptable e-value was set to 10^−10^. We retained the best high-scoring alignment of each query-reference pair based on the highest bit score. We removed data contained in negative controls, alongside ASVs covered by fewer than 15 reads (see SI).

To investigate how well the literature- and OBIS-derived biodiversity information were resolved by eDNA and BRUV, we checked the concordance of all data sources on order, family, genus and on species levels. To judge sampling effort and total species diversity based on BRUV and eDNA observations, we inspected species accumulation curves and calculated Good Turing estimators (giving the number of all species based on species already seen in a small sample; Good, 1953), and then compared those values to combined Te Wahipounamu literature and OBIS species records.

To verify the credibility of eDNA information, we checked all eDNA species lists against a comprehensive list of all New Zealand fish (Roberts et al., 2019) and evaluated species assignment and alignment qualities.

To investigate how useful BRUV, eDNA and OBIS records are in detecting regional differences between fish biodiversity, we used Analysis of Similarity (ANOSIM; Clarke 1993). Thereby analysing Jaccard distances (Jaccard, 1912), we looked for significant differences in taxon (species, genus, family, order) overlap depending on various factor combinations, hence checking whether a particular observation method fared better in detecting taxon composition differences either between different field work areas (WJ, FF, LS), or according to protection status (MR or CNTRL).

## Results

Each of our four data sources yielded different species counts. In total, we yielded 116 species (106 Actinopterygii, 10 Chondrichthyes), comprised of 59 species from previously published Te Wahipounamu works, 44 unique species derived from eDNA, 25 from BRUV and 25 from OBIS (large area; see Table 1, Fig. 2, SI. Table 4). While 21 field work sites (Fig. 1a) yielded environmental DNA and BRUV data (Fig. 1b, c) matching local OBIS data could only be obtained for nine field work sites (Fig. 1a, small circles, LS CNTRL, FF, WJ), and hence those finer scaled OBIS data were later excluded from ANOSIM as incomplete data (Fig. 1a, small circles, Fig. 1d). For further comparisons with Good-Turing estimates, we posit the local “real” species count to 68 as the number of unique species observed across literature and OBIS (Fig. 3).

**Table 1:**
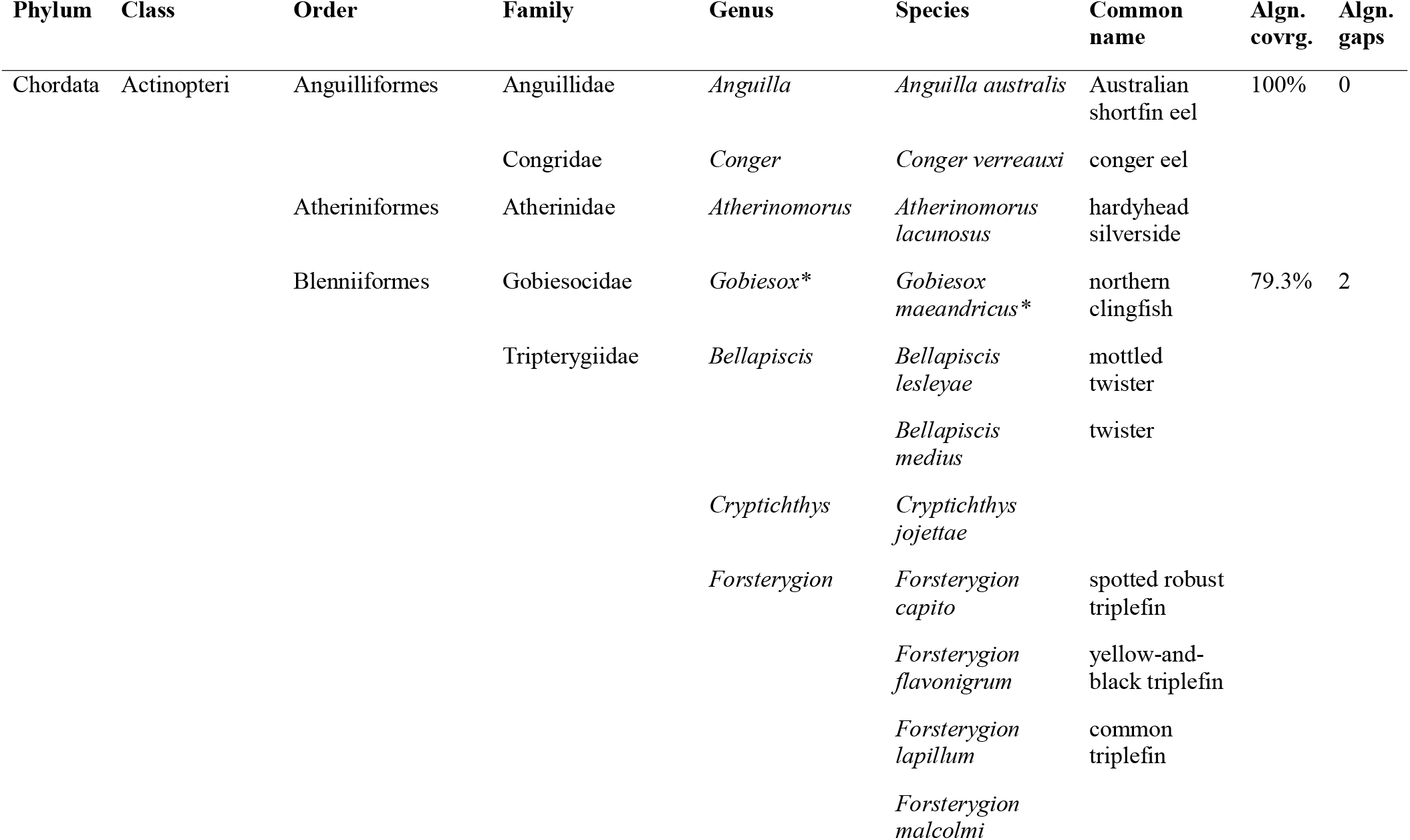
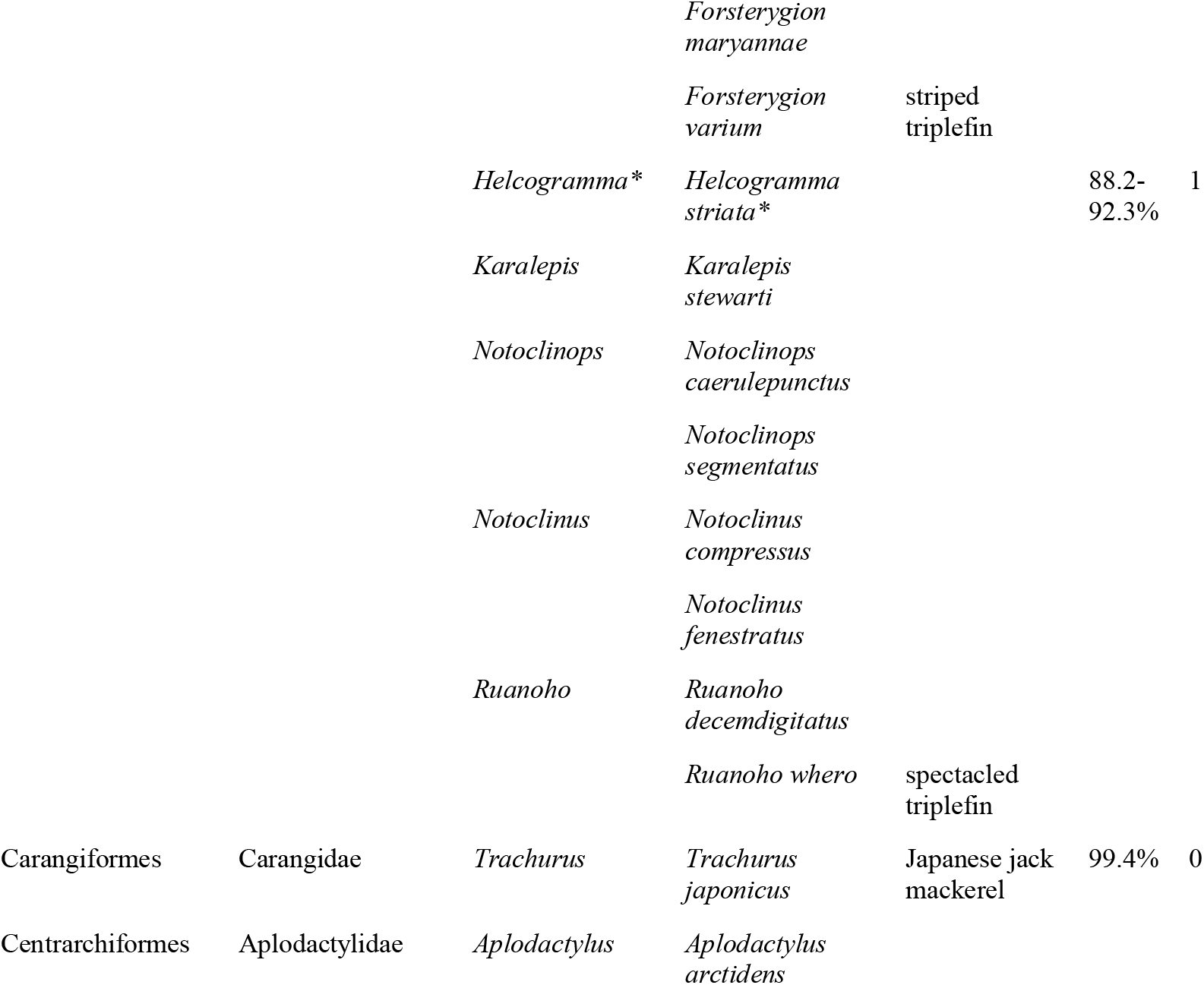
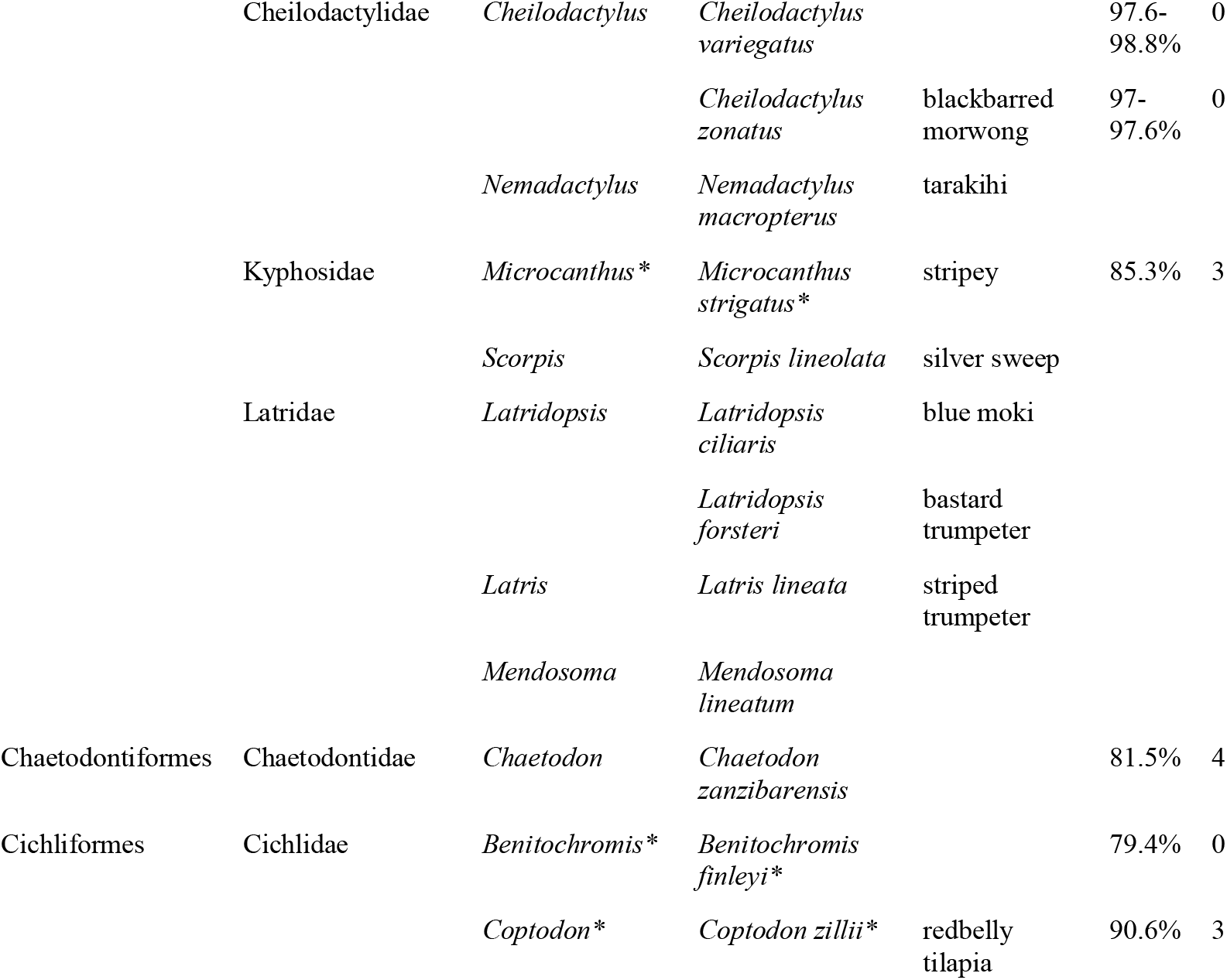
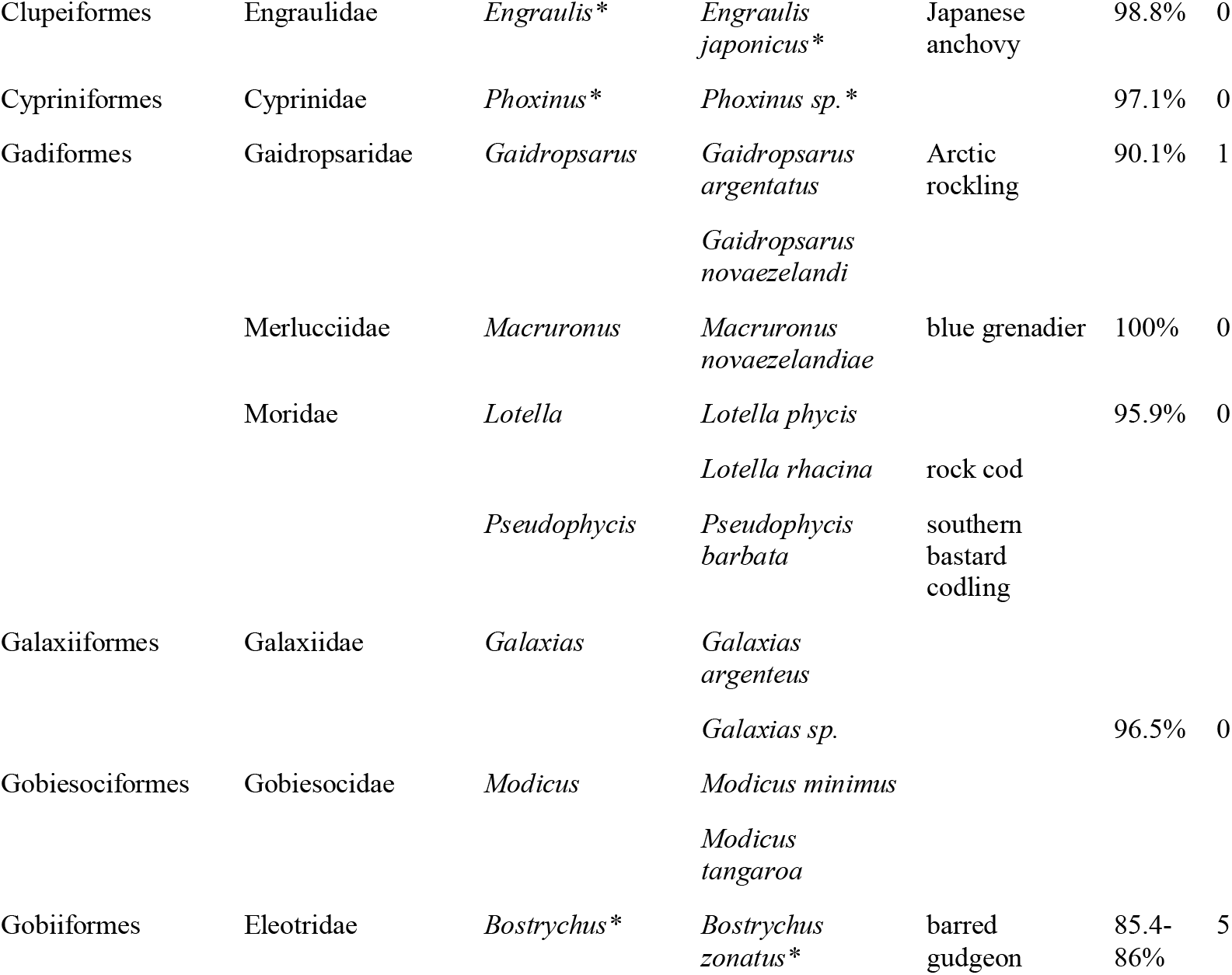
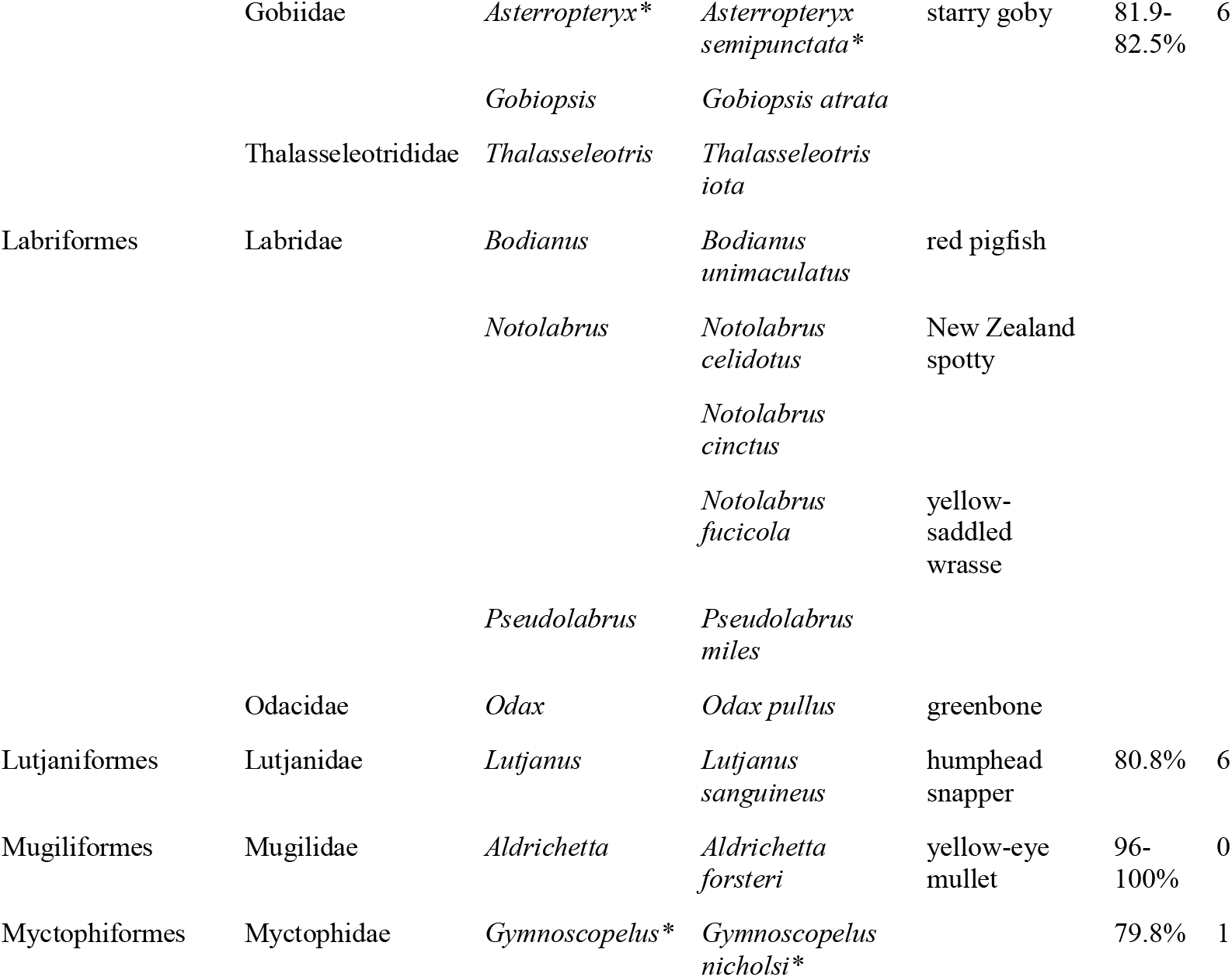
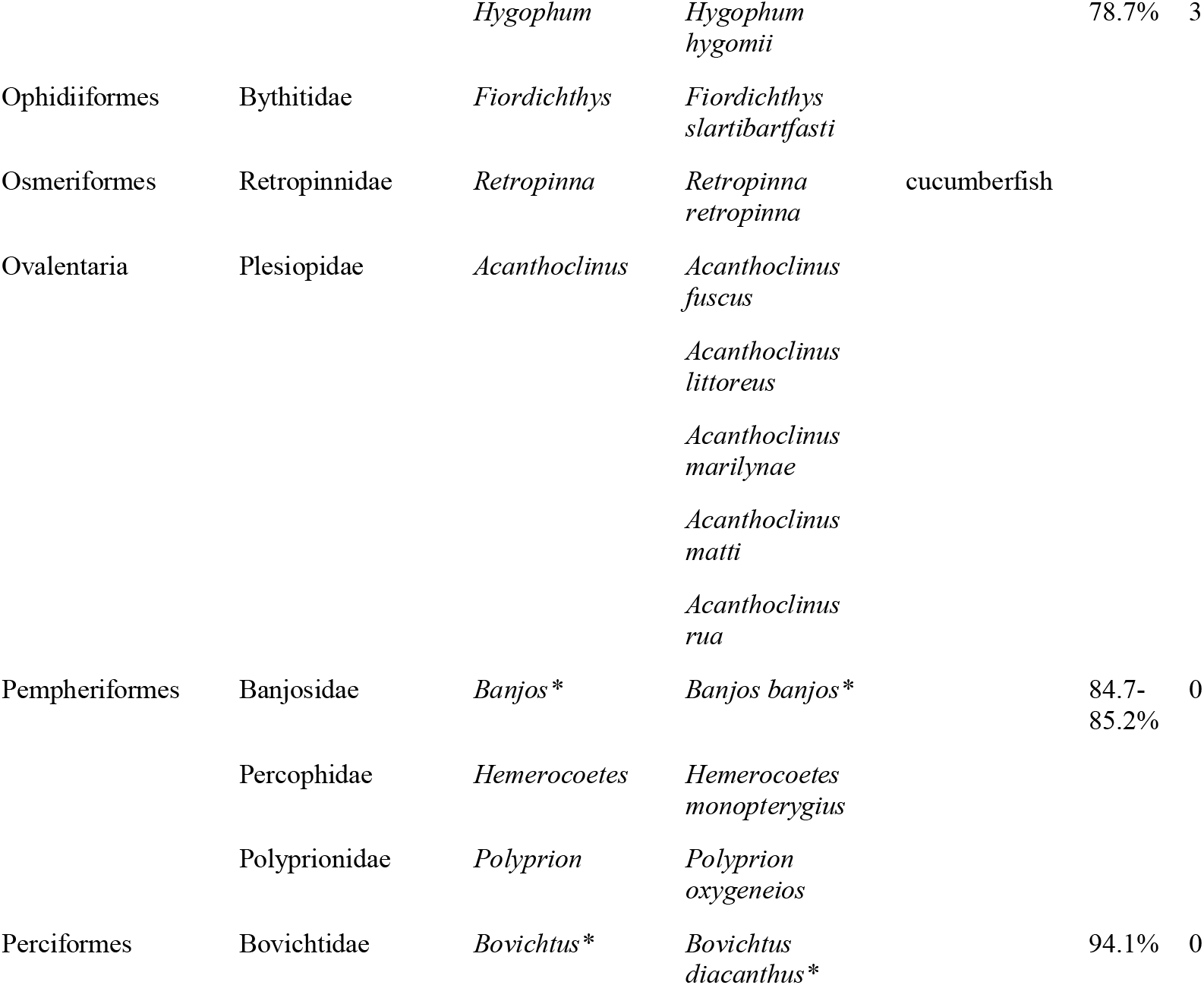
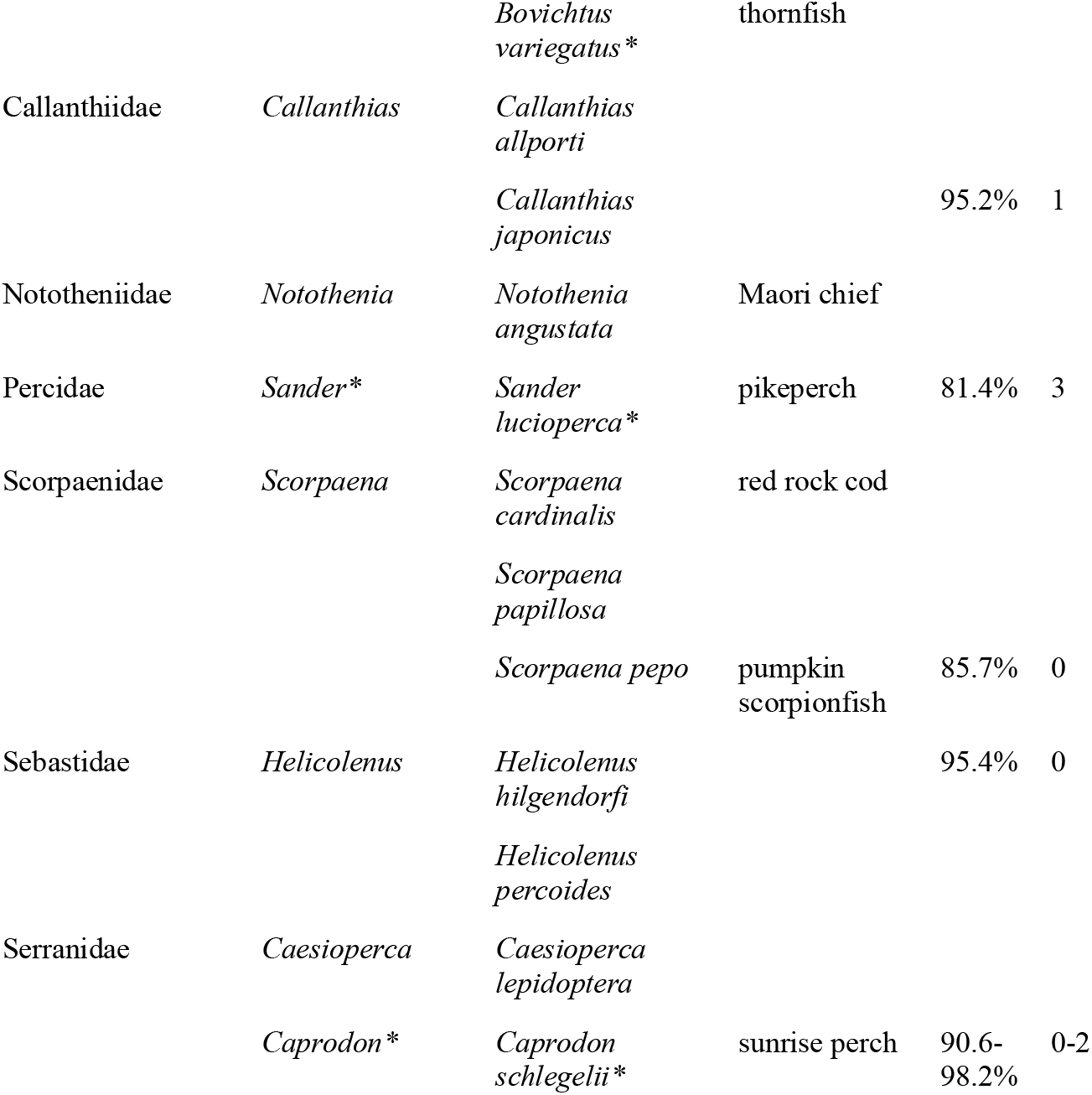
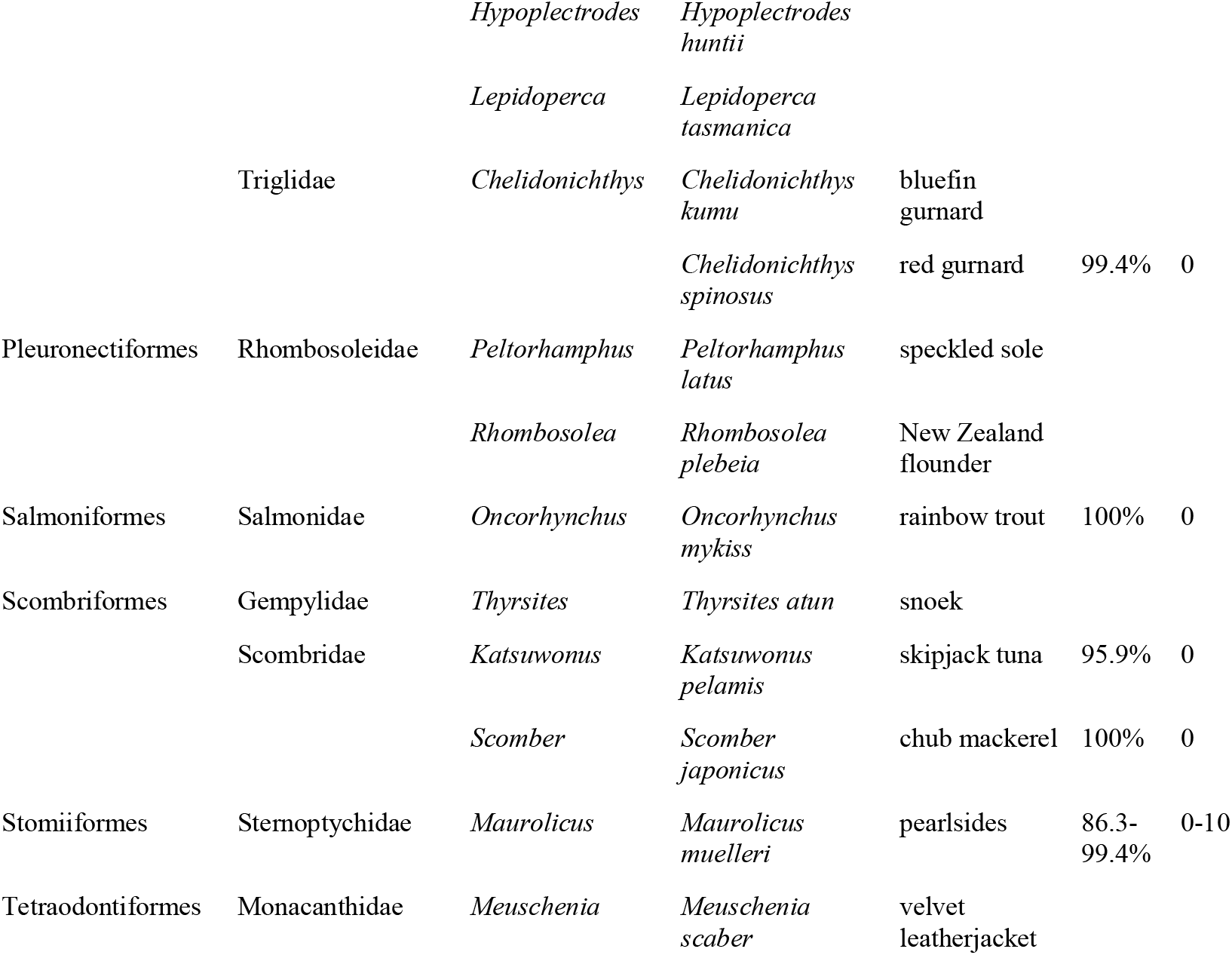
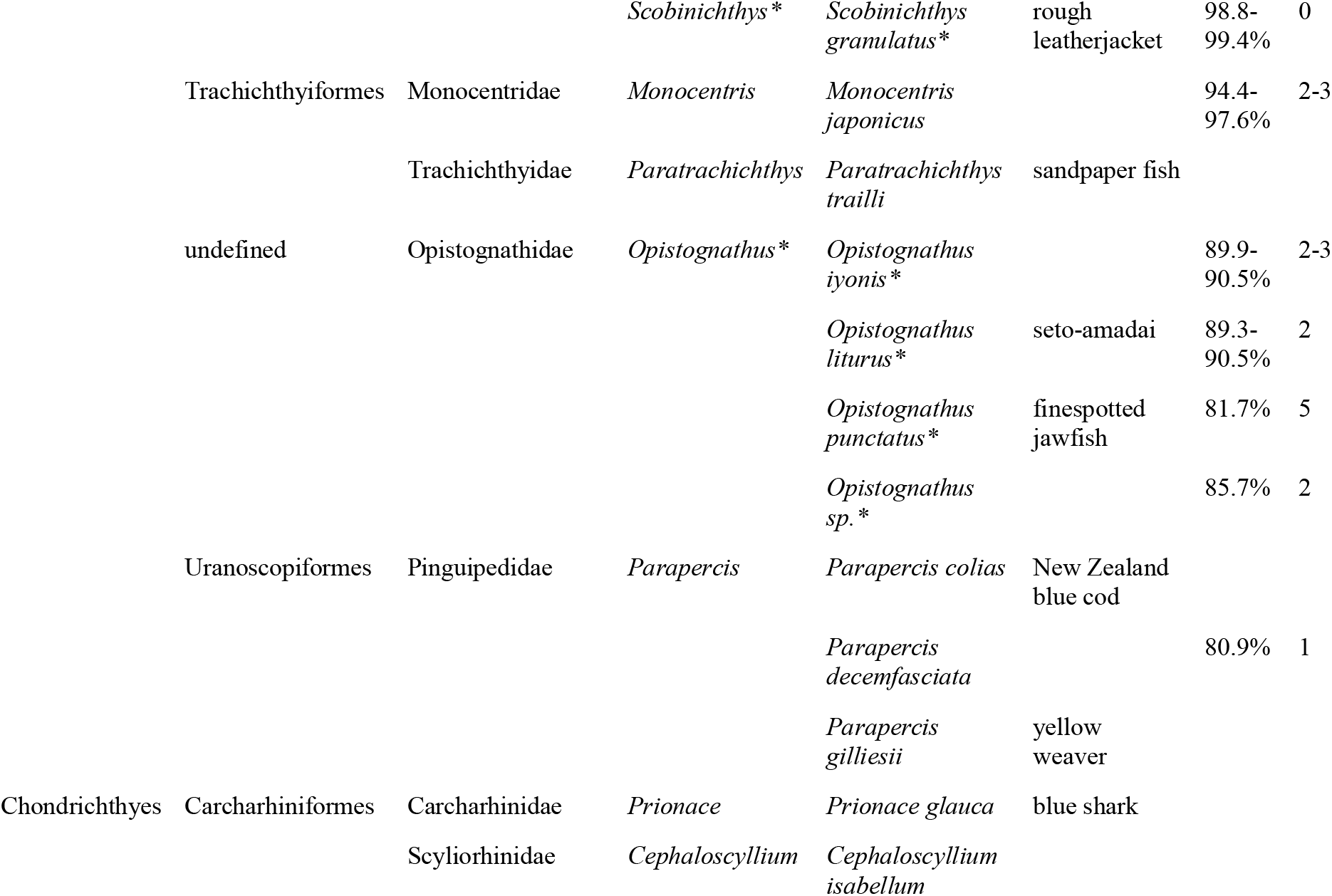
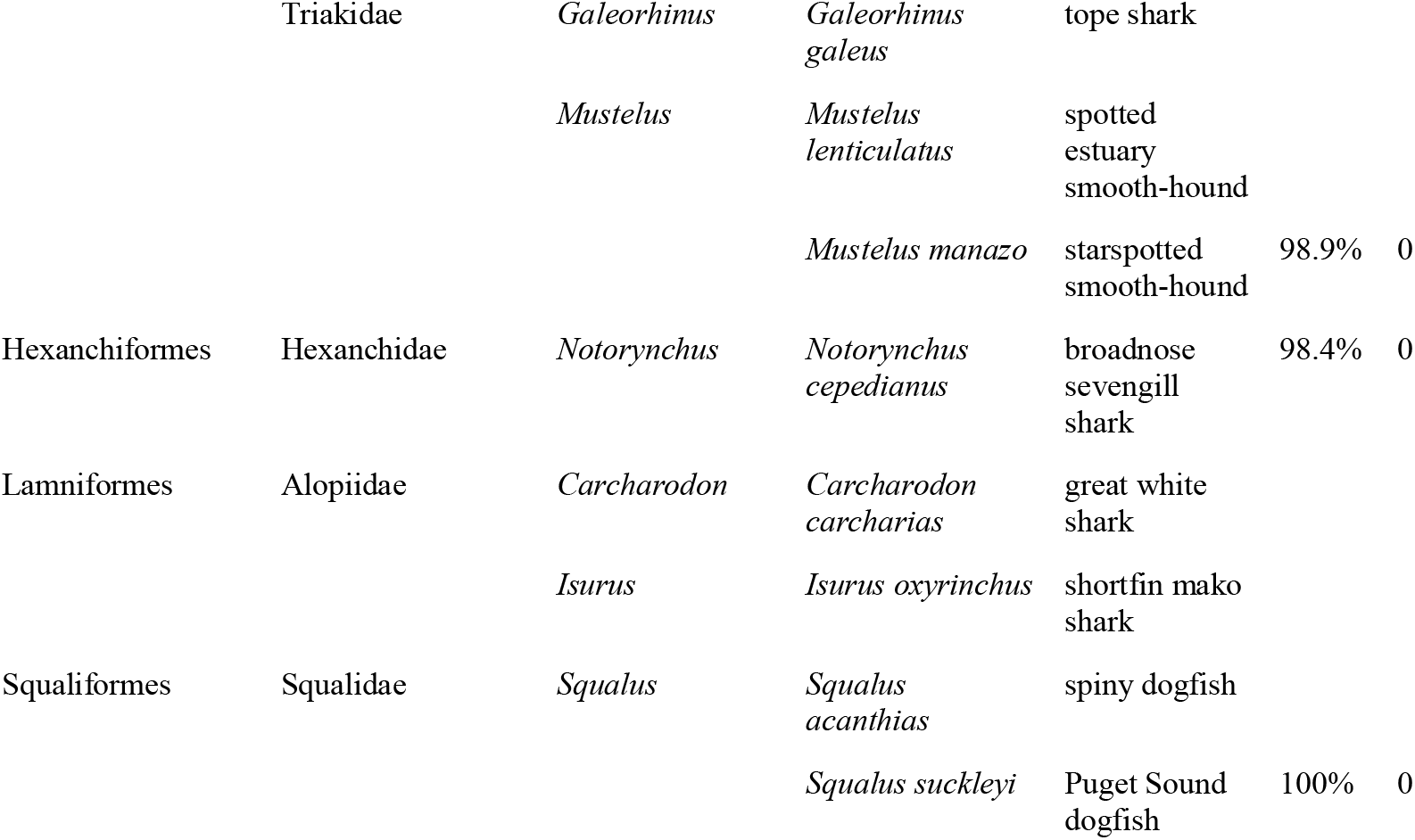
Details on taxonomic observations across data sources. Taxonomic hierarchies conform with NCBI taxonomy where available, thus allow analysis in relation to environmental DNA (eDNA) data and are sorted alphabetically – the resulting species order is identical to Fig. 2. Taxa not listed as New Zealand species by Roberts et al., (2019) are highlighted with asterisk (*). Trivial names are indicated where available from NCBI. For all taxonomic assignments also yield from eDNA we provide the alignment coverage and alignment gaps. Since identical species were assigned to multiple Amplicon Sequence Variants (ASV’s; Callahan et al., 2017) in some instances, ranges are provided for alignment coverages and gap counts for species-specific alignments.

**Fig. 2:**
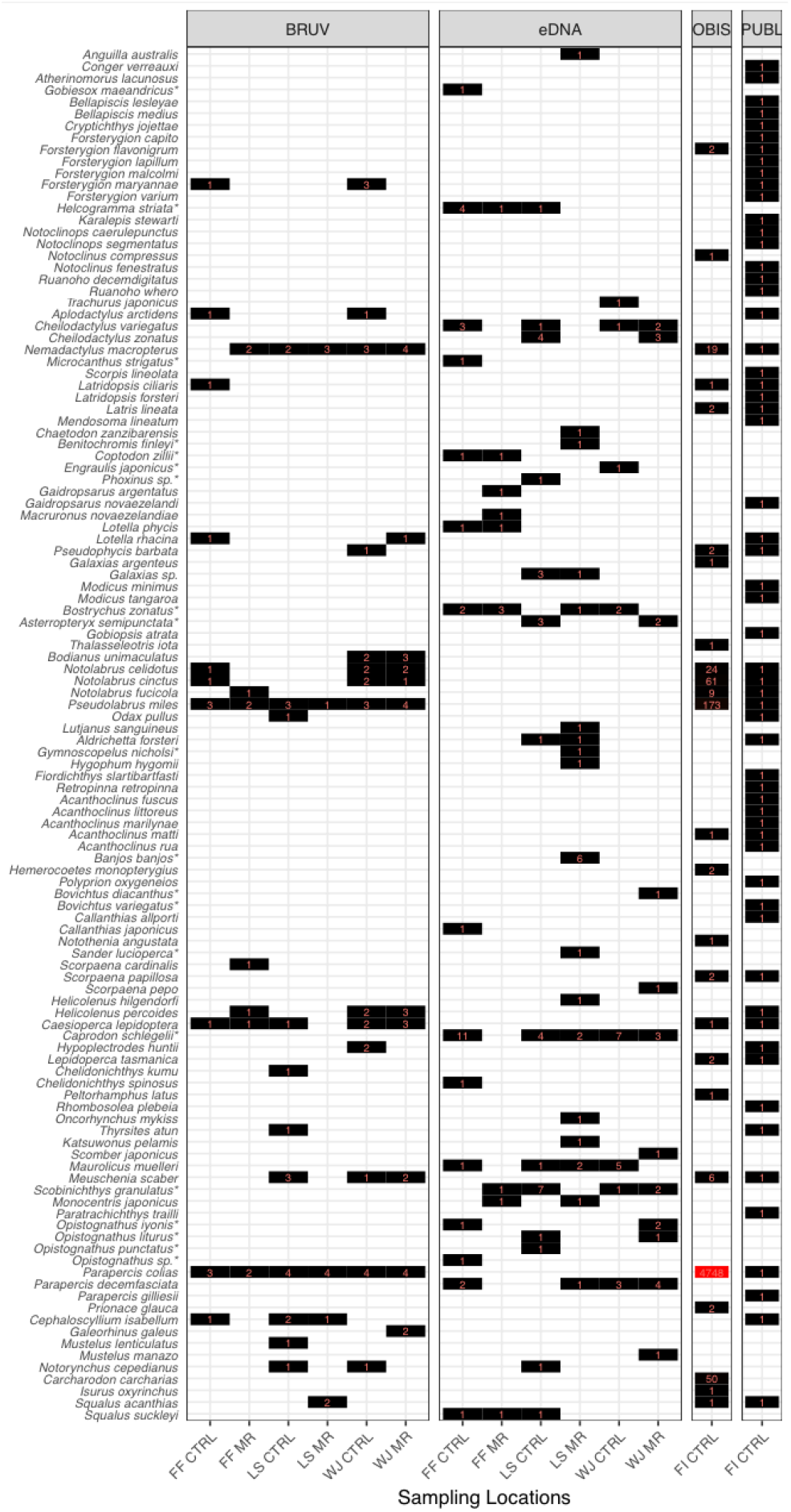
Distinct species observations across data sources and field work locations. Observation types: **BRUV** – Observations from baited remote underwater surveys; **eDNA** – environmental DNA observations; **OBIS** – data retrieved from the Ocean Biodiversity Information System (https://obis.org/) for the area surrounding field work sites (large circle in Fig.1); **PUBL** – Fiordland fish species collated from multiple literature records as summarized by Inglis (2008). Sampling Locations: FF – Five Fingers area; LS – Long Sound area; WJ – Wet Jacket area; MR – marine reserve or commercial exclusion zone; CTRL – neither marine reserve nor commercial exclusion zone. Species list: Order follows Table 1, species not listed as New Zealand Species in (CD Roberts et al., 2019) are marked with an asterisk (*). Graph created using R package *ggplot2 (3.3.5)*.

**Fig. 3:**
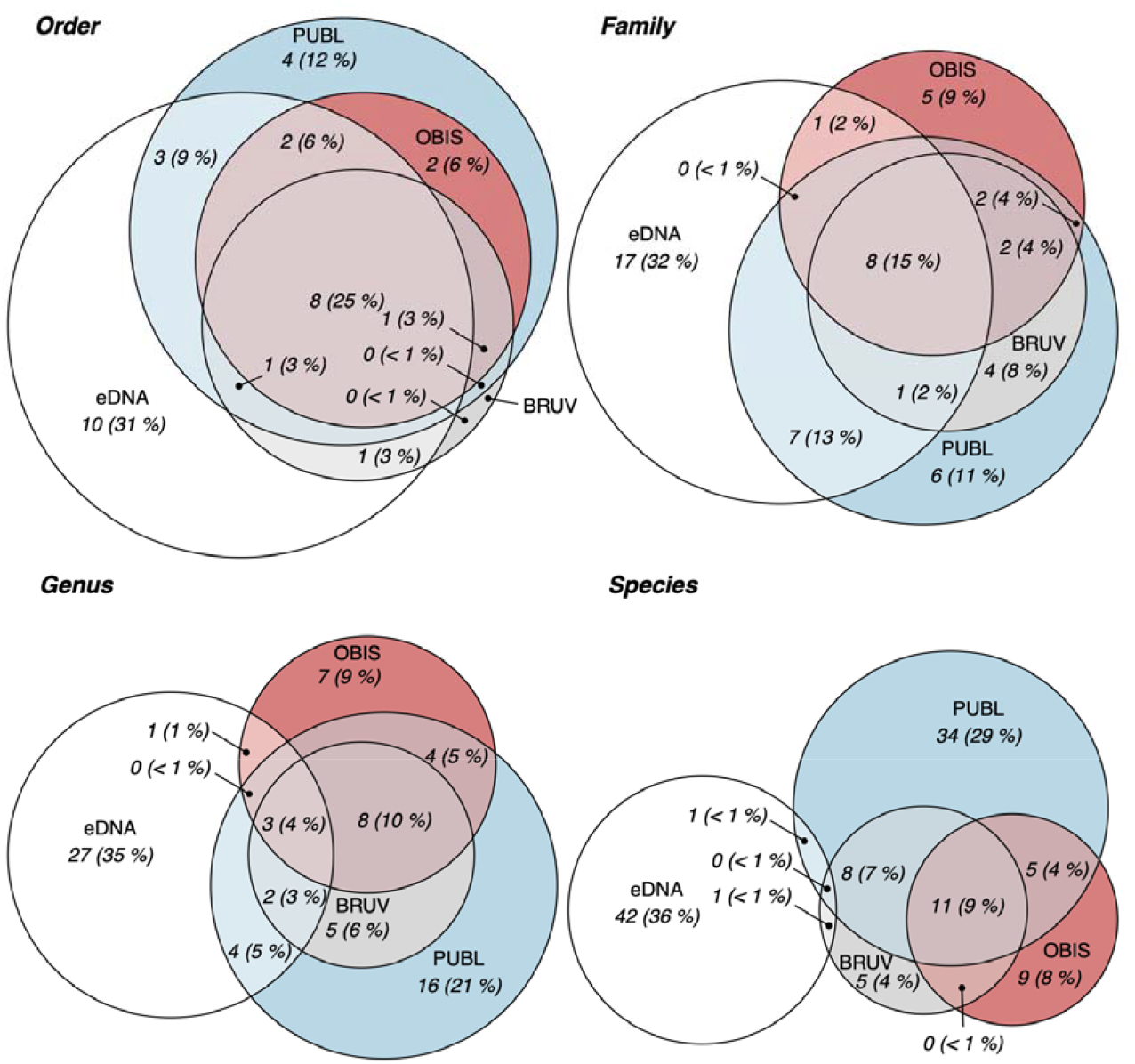
Concordance of taxonomic information across four data sources of Fiordland fish biodiversity. Biodiversity data (Table 1, SI Table 2) is summarized at four different taxonomic levels, shown are unique observation counts at each level, as well as the corresponding percentage of those counts in comparison to all data. Circle sizes proportional to observation count. Observation types: BRUV (grey) – Observations from baited remote underwater surveys; eDNA (white) – environmental DNA observations; OBIS (red) – data retrieved from the Ocean Biodiversity Information System (https://obis.org/) for the area surrounding field work sites (large circle in Fig.1); PUBL (blue) – Fiordland fish species collated from multiple literature records as summarized by (Inglis et al., 2008). Graph created using R package *eulerr (6.1.0)*.

Obtaining community composition comparable to literature and OBIS data within our works’ spatial constraints worked better with BRUV than with eDNA. Nineteen out of 25 species detected with BRUV (76%) were contained in the literature or on OBIS, but only one out of 44 species detected with eDNA (2%) were contained in Te Wahipounamu-specific literature or OBIS (Fig. 1a, large circle). Concordance of taxonomic information between the four data sources dissolved with lowering taxonomic levels, and most decisively for eDNA data (Fig. 3). At species level, only two taxonomic assignments from eDNA matched other data sources, namely *Notorynchus cepedianus* (broadnose sevengill shark), also found with BRUV, and *Aldrichetta forsteri* (yellow-eye mullet) also listed in the literature (Fig. 2). BRUV agreed better with available local biodiversity information, with 11 species mentioned both in the literature and OBIS, and eight detected species mentioned in the literature only (Fig. 3). On BRUV we identified six species (*Bodianus unimaculatus, Chelidonichthys kumu, Galeorhinus galeus, Mustelus lenticulatus, Notorynchus cepedianus, Scorpaena cardinalis*) not mentioned in literature, and not in OBIS, but occurring in New Zealand waters (Roberts et al., 2019). While our plateauing species accumulation curves suggested exhaustive sampling (SI Fig. 5), Good-Turing estimates of eDNA data inferred a presence of 60 species in the study area (assuming 27% missed after 44 observations), and a presence of 26 species using BRUV (assuming 7% missed after 25 observations).

Alignment qualities associated with taxonomic annotation of eDNA data were variable. Forty-four species assigned among eDNA were defined by 92 ASVs (across 142 observations) of which only six yielded flawless alignments with reference data (i.e. 14%, with full query coverage, no alignment gaps). Eighty-six ASVs had variable query coverage (37 families, Tab. 1 and SI), while 32 ASVs had variable gap counts (15 families, Tab. 1 and SI). Mean query coverage was 93.2% (min: 78.6%, med: 97%, sd 6.5%), mean gap count was 1 (max: 10, med: 0, sd 1.83; Tab. 1). Nineteen species assignments among eDNA (43%) had not been observed in New Zealand, and none of these species were found using BRUV, across literature, or OBIS data (apart from *Bovichtus variegatus* – thornfish, not in Roberts et al., 2019, but in Roberts, 2005; Fig. 2). Importantly, 25 species observed with eDNA (56.8% of eDNA-observed species) were known from somewhere New Zealand (Roberts et al., 2019) but were not observed in BRUV or found in Te Wahipounamu literature. Interestingly, using eDNA, we obtained perfect alignments between few ASVs and reference data for *Arctocephalus forsteri* (New Zealand fur seal), *Balaenoptera musculus* (blue whale), and *Tursiops truncatus* (bottlenose dolphin).

In ANOSIM, only BRUV data, and not eDNA data, exhibited location-specific differences among species’ presence overlaps among the 21 sites – on species, genus, family, and order levels. Significant differences were calculated in overlaps between the six field work areas but not between marine reserve nor control areas (SI Table 3).

Investigation of the strikingly homogenous structure of eDNA data by regression analysis of the 142 non-unique eDNA observations (Tjur’s R^2^ 0.027) suggested each additional alignment gap to be associated with a 39% increased probability of observing a non-native species (Odds Ratio 1.39, 95% CI from 1.19 to 1.66, *p* <0.01). A 1% increase in alignment concordance was associated with a 7% increased probability of non-native observation (OR 1.07, 95% CI 1.03–1.12, *p* <0.01). Null deviance was 572.60 on 141 degrees of freedom, residual deviance was 552.31 on 139 degrees of freedom (SI Figs 7 and 8).

## Discussion

What is a realistic estimate of the fish biodiversity in Te Wahipounamu? Based on literature and OBIS alone, we estimate the currently described combined ray-finned and cartilaginous fish species count of Te Wahipounamu to be 68, minding that we constrained OBIS data to surround field sites (Fig. 1, large circle), and that those data are predominantly based on visual observations (SI Table 2). If species counts obtained from literature and OBIS were close to a real value of 68, and the same was true for eDNA and BRUV data, both respective Good-Turing estimates would be 68. Our BRUV-based Good-Turing estimate of 26 species diverges strongly from this number. This may have several reasons. Firstly, we only inspected an isolated area in Te Wahipounamu, while the literature describes a larger area. Secondly, bait in BRUV does not attract all fish for the camera, particularly if deployed at limited depth range, as done here. For eDNA, the Good-Turing estimate of 60 species is more like the literature-inferred species count, but this could be coincidental.

How credible are eDNA derived species assignments with currently available reference data? We believe lacking eDNA reference data to restrict accurate species annotation of ASVs. There are several observations from our data that appear to support this hypothesis. First, while there is a reasonably good concordance between species identified in our BRUV analyses and species known from the area as combined from publications and OBIS, the dissimilarity between eDNA data on one side, and BRUV, OBIS and publication data on the other side, increases with decreasing taxonomic level, culminating in only two out of 44 eDNA species being either identified in our BRUV analyses or known from previous publications (Fig. 3).

Secondly, every approach to identify species diversity in a marine ecosystem has its biases, and published observations are mostly based on visual approaches. Thus, one could argue for the existence of a bias favouring similarity between our visual BRUV observations and published species occurrences to the detriment of eDNA data’s similarity. However, we do not believe this circumstance alone to be responsible for a bias favouring BRUV data to be more similar with literature and OBIS observations in comparison to eDNA observations. Literature and OBIS observation methodologies extend well beyond the specific biases of BRUV, including a multitude of different observation techniques (poison stations, seine net fishing, spear fishing, diver surveys and others, SI Table 1). Collectively, all observation techniques should have provided an appropriately comprehensive overview of fish diversity in Te Wahipounamu, lacking biases inherent to BRUV.

Thirdly, some divergence between eDNA data and the other data sources may be explained by the known ability of eDNA to detect “cryptic” species that are not easily discovered by any visual surveying. The most obvious candidates for this category would be the 25 eDNA species that had previously been reported from New Zealand but not yet from Te Wahipounamu (Fig. 2). However, such a bias should not prevent a broad overlap between eDNA and visual approaches for species that can easily be detected visually. Clearly, we did not find such an overlap.

Crucially, of the 25 species we detected by BRUV and that were therefore present at the time of our concurrent water sampling for eDNA analyses, 24 species are not present in the NCBI reference database (Fig. 2) and could therefore not be detected by our eDNA approach. This highlights one of the main limitations of eDNA multispecies surveys today.

Nevertheless, and despite the lack of reference data, eDNA still identified a larger number of species than our concurrent BRUV analyses. From where do these species assignments come? In most cases during taxonomic assignment, where no perfect match can be found between eDNA query and reference subject sequence, the algorithm assigning ASVs to species information (BLAST) chose the next-closest matching species contained in the reference data collection, as encouraged by our taxonomic assignment parameters. Our taxonomic assignments correspond with this hypothesis, as binomial regression showed that each additional gap in a sequences’ reference alignment associated with a 39% increased probability of observing of a non-native species.

Interestingly, a 1% increase in alignment concordance increased the likelihood of a non-native observation as well, by 7%. At first sight this seems counter-intuitive, however the latter observation is also consistent with our hypothesis: A poorly matching sequence would not be assigned to a matching species but rather to a higher matching taxon such as genus or family. A better fit increases the likelihood of a species level assignment, but without native species contained among reference data, the likelihood increases that the query sequence is assigned to a closely related species not occurring in New Zealand. Similar observations have been made in other regions of the world (Stoeckle, Das Mishu, & Charlop-Powers, 2020).

The large number of species detected by our eDNA approach – although probably misassigned in several instances – is a testament to the potential power of eDNA methods. Arguably, any detected effect of lacking reference data could be less pronounced by using another, or multiple primer pairs. For example, our primer evaluations with the recently released software GAPeDNA (Marques et al., 2021) show that the “Fish 16S” primer set by McInnes et al. (2017) would have covered 249 instead of the 119 New Zealand marine fish species covered by our MiFish 12S dataset (SI Table 1). However, the overall conclusion remains. Of the over 1294 known New Zealand marine fish species, molecular reference data of any kind is available only for 489 species in southern New Zealand, and for no available primer pairs sufficient reference data is available. Hence without substantial effort into generating suitable reference data for a carefully selected range of similar primers, eDNA analysis here and everywhere else will remain an impaired tool for biodiversity management. While this insight holds true after almost two decades of eDNA research (Hebert, Cywinska, Ball, & DeWaard, 2003) we note that a growing number of researchers are working hard on closing reference data voids around the globe (reviewed in Marques et al., 2021; Weigand et al., 2019).

## Acknowledgments and Data

Funded in part by the New Zealand Department of Conservation. Authors were supported by the University of Otago. Authorship determination followed the Contributor Roles Taxonomy (https://casrai.org/credit). P.C. lead field, lab and analysis work and wrote manuscript. M.d.L. provided Good-Turing estimates, accumulation curves and advised P.C. M.H., M.L., P.C. obtained, and M.H. viewed BRUV footage. M.H. aided DNA extraction. W.R., C.H., M.K. obtained funding. Everyone contributed to manuscript revision. We appreciate Olga Kardailsky for obtaining deceased aquarium fish, and Anya Kardailsky for helping water filtering (both University of Otago). Sequencing was undertaken by Otago Genomics. Data and code generated for this work are archived at https://doi.org/10.5281/zenodo.4638297.

