## Supplementary material for "Environmental DNA analysis needs local reference data to inform taxonomy-based conservation policy – A case study from Aotearoa / New Zealand": SI

October 22, 2021

### Contents

**Contents** 1

**List of Figures** 2

**List of Tables** 2

**1 Materials and methods** 2

**References** **11**

**A Taxonomic observations** **16**

### 1 Materials and methods

#### Data collation and analysis

All analyses were conducted in R (R Core Development Team, 2019) and Megan (Huson et al., 2016) of increasing versions. Packages used for analysis in R included *curl, data.table, decontam, dplyr, flextable, future.apply, ggpubr, grid, gridExtra, indicspecies, janitor, knitr, magick, magrittr, officer, openxlsx, phyloseq, primerTree, readxl, reshape2, robis, sf, sjPlot, sp, taxize, taxonomizr, tidyr, tidyverse, vegan* and *xaringan* (Auguie, 2017; Bache and Wickham, 2020; Bengtsson, 2021; B. Callahan and Davis, 2021; Chamberlain et al., 2020; Dowle and Srinivasan, 2021; Firke, 2021; Gohel, 2021a; Gohel, 2021b; Hester, 2020; Kassambara, 2020; Lüdecke, 2021; McMurdie et al., 2021; Oksanen et al., 2015; Ooms, 2021a; Ooms, 2021b; Pebesma, 2021; Pebesma and Bivand, 2021; Provoost and Bosch, 2021; Schauberger and Walker, 2021; Sherrill-Mix, 2021; Wickham, 2020; Wickham, 2021a; Wickham, 2021b; Wickham and Bryan, 2019; Wickham, François, et al., 2021; Xie, 2021a; Xie, 2021b).

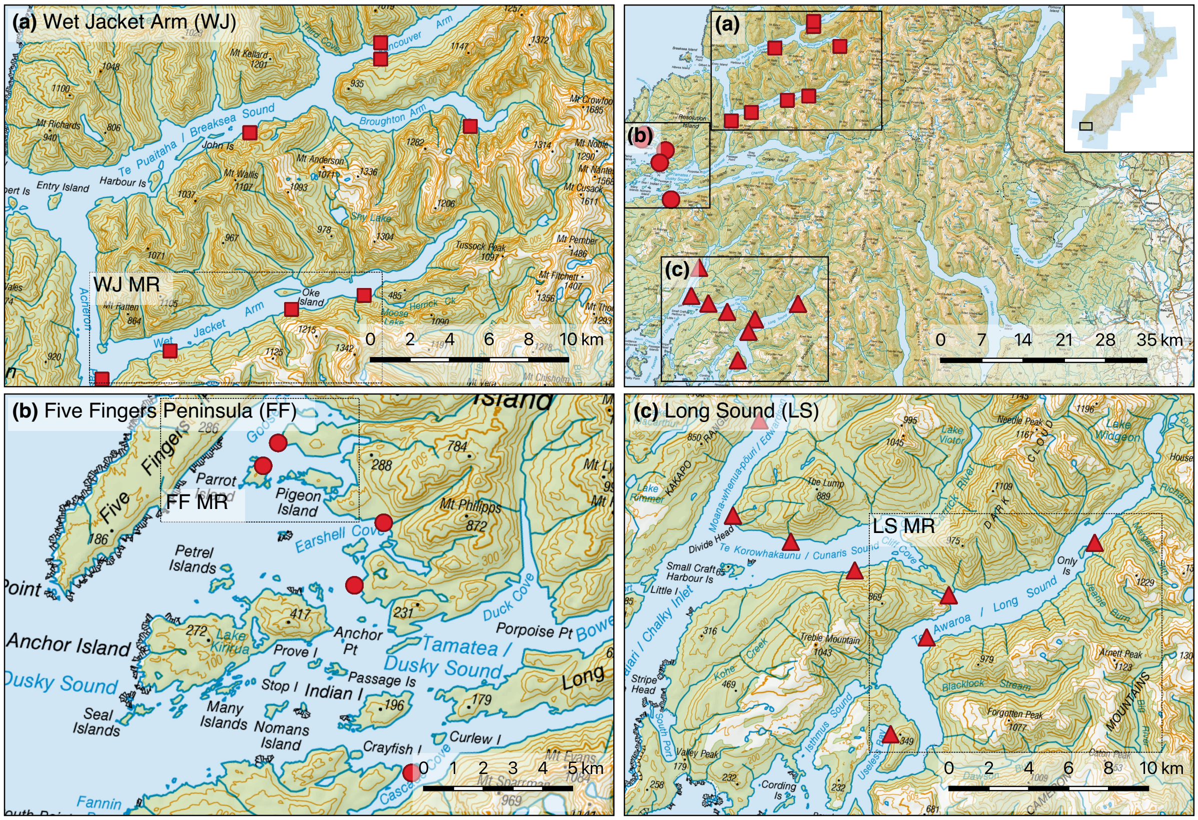

**Figure 1:** Sampling locations for BRUV and eDNA data collection in Te Wahipounamu, New Zealand. We analyzed species observations from 21 locations. Within each of the three sampling regions (WJ, FF, and LS), samples collected inside MRs are delineated by hatched rectangles, control samples are outside of rectangles. Base layer sourced from the LINZ Data Service licensed for reuse under CC BY 4.0.

### Field work

To obtain eDNA at each sampling location (Fig.1), duplicates of 900 ml water portions were collected using a bleached Niskin Bottle, passed through separate 0.22 µm Sterivex columns (Merck, US-NJ) using sterile disposable 50 ml syringes, akin to Jeunen et al. (2020). To control contamination, negative controls were obtained after every second sample by passing molecular grade distilled water through third columns. After filtering, columns were filled with 5 ml Longmire's solution, sealed, and stored at 4°C for 21 days and -20°C subsequently (Majaneva et al., 2018).

BRUV footage was obtained for one hour using two waterproof cameras, recording at 1080p resolution, at 60 frames per second, 40 cm apart, at a converging angle of 8°, aided by two 300 lm lights. We filmed a permeable plastic container suspended 1.2 m below the cameras, carrying 500g bait *(Sardinops sagax)*, akin to (Jeunen et al., 2020). Video footage were analyses by eye, taxonomic observations were formalized for further analysis using the NCBI taxonomy database (Federhen, 2012) and R package *taxonomizr* (Sherrill-Mix, 2021).

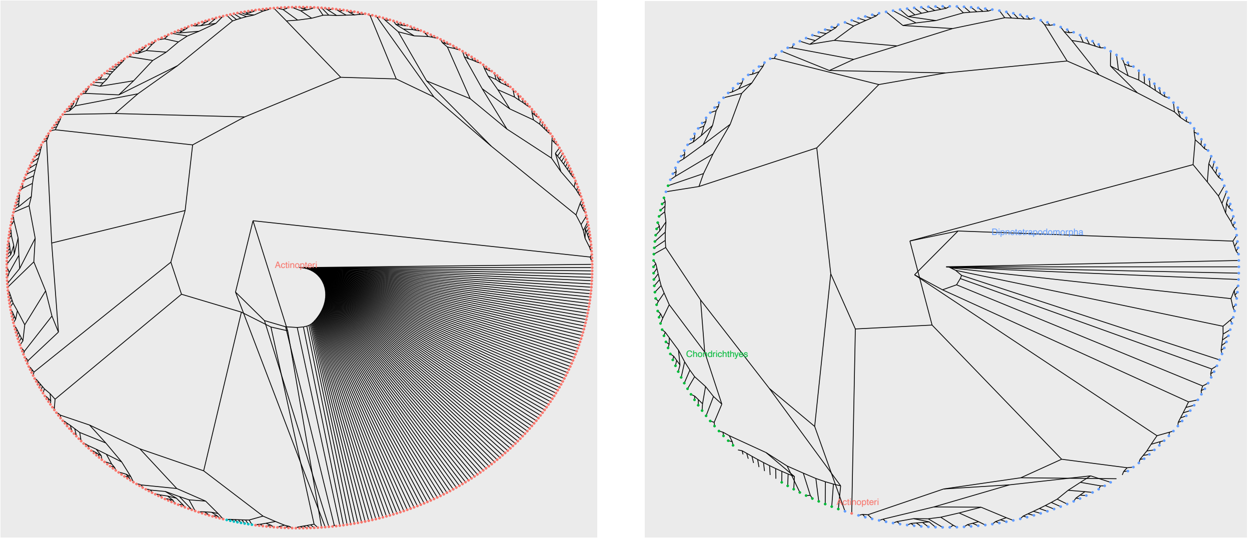

**(a)** Mifish U: ray-finned fish, tetrapods and lungfish.

**(b)** Elas02: cartilaginous and ray-finned fish, tetrapods, and lungfish

**Figure 2:** *In silico* PCR of used primer pairs: Likely amplification specificity of Mifish U and Elas02.

#### Primer selection and *in silico* PCR

Primer selection was verified using GAPeDNA v.1.0.1 (https://shiny.cefe.cnrs.fr/GAPeDNA/; Marques et al. 2021) and Scite (Nicholson et al., 2021). We assed our primer selection by initially evaluating online information provided by GAPeDNA with regards to reference data coverage for marine fish in the southern New Zealand province for all primer pairs listed. Subsequently, we retrieved the number of publications referencing the work introducing the primers.

*In silico* PCR of primers “MiFish-U” (Miya et al., 2015) and “Elas02” (Taberlet et al., 2018) was performed with R package *primerTree* (Hester, 2020) with default parameters (500 alignments, three possible mismatches).

#### Environmental DNA data processing

Amplicon Sequence Variants (ASV’s *sensu* B. J. Callahan, McMurdie, and Holmes 2017) were generated from raw sequence data including use of Qiime v2 2020-08 (Bolyen et al., 2019). Initially, sequence quality was checked with FastQC (Andrews, 2010) as called by MultiQC (Ewels et al., 2016). We then used Cutadapt v3.0 (Martin, 2011) to deconvolute samples, while not allowing for any mismatches, and choosing an Expected Error value (Edgar and Flyvbjerg, 2015) of zero, to obtain high quality data.

Deconvoluted data were imported into Qiime and denoised (Rosen et al., 2012) using the DADA2 algorithm (v1.10.0; B. J. Callahan, McMurdie, Rosen, et al. 2016). Taxonomic annotation of denoised sequence data was obtained using Blast 2.10.0+ (Camacho et al., 2009) and a recent copy (April 2020) of the NCBI nucleotide collection (Benson et al., 2011) excluding environmental samples, while requiring a minimum identity 75%, a minimum query coverage of 95% and an e-value of $10^{-10}$, retaining the first high-scoring alignment of each query-subject pair based on Bit-score (see Fig. 3 on the next page for raw data summary). To mitigate impact

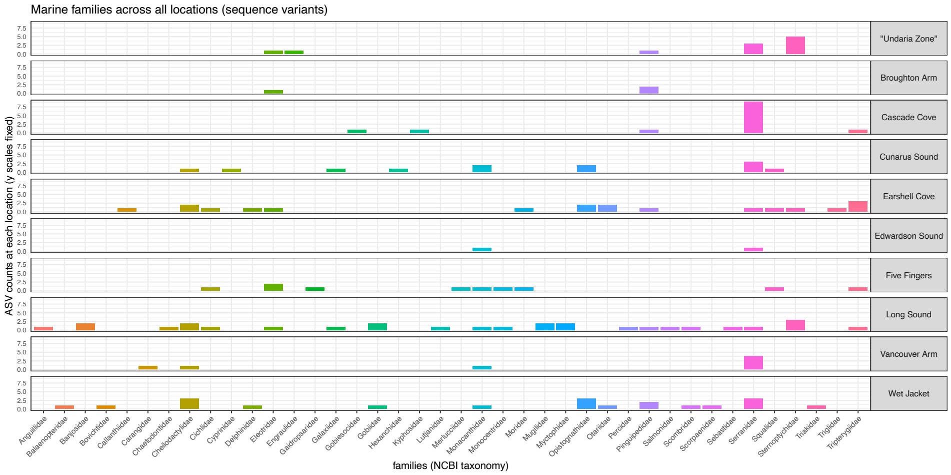

**Figure 3:** Taxonomic assignments for all sample types among raw data, prior to filtering. `blank`: field water controls, `eDNA`: study observations, `nctrl-pcr`: PCR negative controls, `nctrl-xtr`: Extraction negative controls, `pcntrl-blnd`: positive controls.

of minor cross-contamination or tag jumping in our data (Schnell et al., 2015), as detected using our positive controls, we analyzed and subtracted negative control data from all samples before statistical analysis, which included using R package *decontam* (B. Callahan and Davis, 2021), and removing ASVs with covered by less the 15 reads each.

For statistical analysis, taxonomic observations from eDNA were formalized with R packages *taxize* and *taxonomizr* (Chamberlain et al., 2020; Sherrill-Mix, 2021) akin to BRUV, OBIS, and literature data. After inspection of all eDNA and BRUV derived species assignments, eDNA species lists were limited to classes Actinopteri and Chondrichthyes, and all other data was deemed irrelevant for comparison with BRUV observations (crustaceans, mollusks, and hagfish seen on BRUV were absent from eDNA data prior to data filtering, keeping in mind the amplification profile shown in Fig. 2 on the preceding page).

#### Indicator species analysis

To find species that could be used as ecological indicators of associated ANOSIM-distinguished community or habitat types, we used indicator species analysis (De Cáceres et al., 2010) on the data set used for ANOSIM.

#### Binomial regression

To further test the credibility of eDNA information test and whether incompletely resolved ("cryptic") eDNA ASVs inflated the number of species observed across multiple locations during ANOSIM, we used binomial regression. The response variable was non-native status of eDNA detected species (CD Roberts et al., 2019). Predictor variables were the number of alignment gaps (integer), and alignment query coverage (percentage). The number of trials was defined using by numbers of locations at which each assigned species was seen.

**Table 1:** Evaluation of primer selection through GAPeDNA v.1.0.1 (https://shiny.cefe.cnrs.fr/GAPeDNA/; Marques et al. 2021). Listed is reference data coverage for marine fish in the southern New Zealand province as provided by the web tool, and citation count of primer references. GAPeDNA lists 489 sequenced species for southern New Zealand (11-Sep-2021). Citation counts obtained 13-Sep-2021. Total distinct citations as estimated by Scite (Nicholson et al., 2021).

| Primer region | primer | species sequenced | reference | citations |
|---|---|---|---|---|
| 12S | ACMDB | 27% (134) | Bylemans et al. 2018 | 45 |
| | Ac12S | 29% (141) | Evans et al. 2016 | 229 |
| | Am12S | 25% (124) | Evans et al. 2016 | 229 |
| | - | 31% (154) | Kelly et al. 2014 | 263 |
| | MiFish-U | 24% (119) | Miya et al. 2015 | 378 |
| | Teleo | 24% (117) | Valentini et al. 2016 | 545 |
| 16S | - | 51% (249) | DiBattista et al. 2017 | 42 |
| | Ac16S | 19% (95) | Evans et al. 2016 | 229 |
| | Ve16S | 36% (177) | Evans et al. 2016 | 229 |
| | - | 29% 144 | Kitano et al. 2007 | 148 |
| | Fish | 51% (249) | McInnes et al. 2017 | 23 |
| | ar / br | 21% (101) | Palumbi 1996 | *undeterminable* |
| | - | 51% (249) | Shaw et al. 2016 | 141 |
| COI | coi1 | 6% (29) | Ivanova et al. 2007 | 877 |
| | vf1d | 16% (80) | Ivanova et al. 2007 | 877 |
| | F1 | 10% (48) | Ward et al. 2005 | 2 327 |
| | F2 | 6% (30) | Ward et al. 2005 | 2 327 |
| CytB | - | 13% (66) | Kocher et al. 1989 | 3 536 |
| | - | 17% (85) | *M. Miya - undeterminable* | *undeterminable* |
| | 2cb | 8% (41) | Thomsen et al. 2012 | 521 |
| | 2de | 15% (74) | Thomsen et al. 2012 | 521 |
| | cb | 7% (36) | Thomsen et al. 2012 | 521 |
| 18S | - | 2% (11) | *J. McDondald - undeterminable* | *undeterminable* |

### 2 Results

#### Primer selection and *in silico* PCR

Results of our primer assessment are shown in Table 1. *In silico* analyses found the employed primers to amplify some tetrapods as shown in Fig. 2 on page 4.

#### Environmental DNA data processing

The denoised, quality-filtered sequenced data consisted of 3 877 007 sequences across 125 samples and 2 139 ASV's (436 Eukaryota, as well as 1 703 non-eukaryote, i.e. Bacteria, Viruses, and undefined taxa; furthermore taxa across 5 super-phyla, 31 phyla, 56 classes, 151 orders, 254 families, 426 genera, and 914 species). Sample mean coverage was 31 016 reads (min.: 1, med.: 2 001, max.: 272 614, standard deviation 54 592), and ASV mean (min, median, max) coverage was 1 812 reads (min.: 1, med.: 18, max.: 803 140, standard deviation 24 270).

Notable contaminants most likely introduced during water filtering (and subsequently removed) included *Homo sapiens* (human, 30 1070 reads), and *Cervus elaphus* (red deer, 19 372 reads). Low abundant ASVs

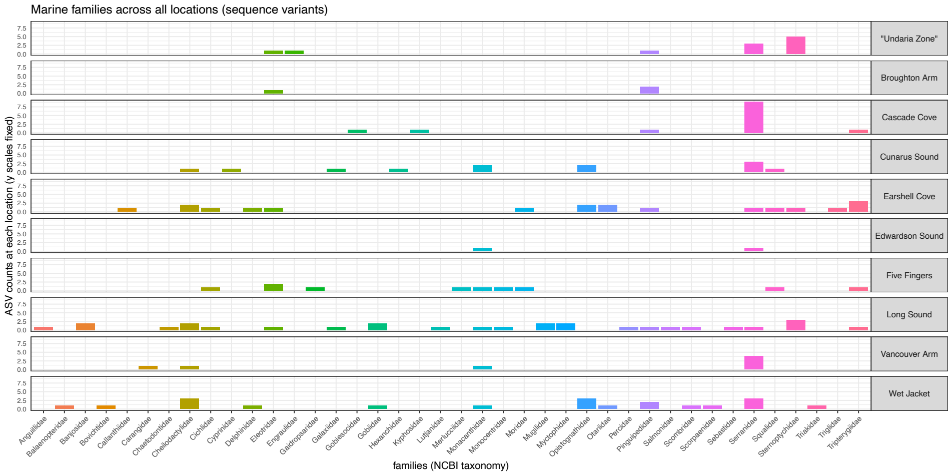

**Figure 4:** Family level assignments and ASV counts among clean data. For location names refer to Fig. 1 on page 3. For species level assignments refer to Table 4 on page 17. For highly highly detailed information refer to the complete study data provided as supplemental online material.

removed as possible cross-contamination not associated with humans included *Trichosurus vulpecula* (bush-tail possum, 16 reads), and *Sus scrofus* (wild boar, 11 reads). We deem the contaminant profile to be very similar to a study conducted in Stewart island (southern New Zealand) with the same primers (Jeunen et al., 2020).

After completed filtering sequence data contained 157 100 sequences across 43 samples and 96 ASV's (96 Eukaryota; including taxa across one super-phylum, 1 phylum, 3 classes, 28 orders, 40 families, 43 genera, and 47 species; also see Fig. 4. Sample mean coverage was 3 353 reads (min.: 32, med.: 1 504, max.: 49 976, standard deviation was 7 940), and ASV mean coverage was 1 636 reads (min.: 17, med.: 304, max.: 24 182, standard deviation 3 793).

Three (ray-finned) fishes not identifiable from video footage had been removed from the final species observation list. NCBI taxonomy did not contain entries for five video-observed species *(Forsterygion maryannae, Cephaloscyllium isabellum, Hypoplectrodes huntii, Notolabrus cinctus, Eptatretus cirrhatus).*

#### Summaries of taxonomic observations

A summary of observations across data sources is listed in Table 4 on page 17. From Grange (1985), Inglis et al. (2008), Mladenov (2001), Clive Roberts (2005), and Wing and Jack (2013) we included into our analysis the species *Acanthoclinus fuscus, Acanthoclinus littoreus, Acanthoclinus marilynae, Acanthoclinus matti, Acanthoclinus rua, Aldrichetta forsteri, Aplidium adamsi, Aplodactylus arctidens, Aplodactylus arctidens, Atherinomorus lacunosus, Bellapiscis lesleyae, Bellapiscis medius, Bovichtus variegatus, Caesioperca lepidoptera, Callanthias allporti, Cephaloscyllium isabellum, Cominella sp., Conger verreauxi, Cryptichthys jojettae, Eptatretus cirrahtus, Fiordichthys slartibartfasti, Forsterygion flavonigrum, Forsterygion lapillum, Forsterygion malcolmi, Forsterygion varium, Gaidropsarus novaezelandi, Modicus minimus, Gobiopsis atrata, Grahamina capito, Helicolenus percoides, Hypoplectrodes huntii, Karalepis stewarti, Latridopsis ciliaris, Latridopsis forsteri, Latris lineata, Lepidoperca tasmanica, Lissocampus filum, Lotella rhacina, Mendosoma lineatum, Modicus tangaroa, Nemadactylus macropterus, Notoclinops caerulepunctus, Notoclinops segmentatus, Notoclinus fenestratus, Notolabrus celidotus, Notolabrus cinctus, Notolabrus fucicola, Forsterygion maryannae,*

**Table 2:** Observation methods across literature sources and OBIS: Species observations listed in CD Roberts et al. (2019) for all of New Zealand were not used as part of the literature corpus analyses of Te Wahipounamu, but only to confirm native status of eDNA assignments.

| Data source | Observation methods |
|---|---|
| Inglis et al. 2008 | literature survey, poison stations and beach seine netting |
| Grange (1985) | unknown |
| Mladenov (2001) | unknown |
| Clive Roberts (2005) | Rotenone ichthyocide, spear gun, fishing with baited lines |
| Wing and Jack (2013) | diving surveys for conspicuous reef fish |
| OBIS | 6 192 human observations, 62 machine observations, 59 preserved specimen |
| CD Roberts et al. (2019)* | voucher specimen described by taxonomic ichthyologist, validated species from literature, clear published account |

*Odax pullus, Parapercis colias, Parapercis gilliesii, Paratrachichthys trailli, Meuschenia scaber, Patiriella regularis, Polyprion oxygeneios, Pseudolabrus miles, Pseudophycis barbata, Retropinna retropinna, Rhombosolea plebeia, Ruanoho decemdigitatus, Ruanoho whero, Scorpaena papillosa, Scorpis lineolata, Squalus acanthias,* and *Thyrsites atun*. Observation methods for these species are listed in Table 2.

Notable taxa observed during environmental DNA (eDNA) processing included[1,2]

- in field controls: *Homo sapiens*♠, *Cervus elaphus*♠, *Serranidae* sp., *Poecilia reticulata*♣, *Pictilabrus laticlavius*$^{*}$
- in positive controls: *Poecilia reticulata*♣, *Chromobotia macracanthus*$^{*}$, *Poecilia latipinna*♣, *Poecilia formosa*♣, *Corydoras aeneus*$^{*}$
- in negative controls: *Homo sapiens*♠, *Pictilabrus laticlavius*$^{*}$, *Poecilia latipinna*♣

Actinopteri obtained through eDNA analysis and video footage included:

- in reserves, but not outside: *Anguilla australis, Banjos banjos, Benitochromis finleyi, Bovichtus diacanthus, Chaetodon zanzibarensis, Gaidropsarus argentatus, Gymnoscopelus nicholsi, Helicolenus hilgendorfi, Hygophum hygomii, Katsuwonus pelamis, Lutjanus sanguineus, Macruronus novaezelandiae, Monocentris japonicus, Notolabrus fucicola, Oncorhynchus mykiss, Sander lucioperca, Scomber japonicus, Scorpaena cardinalis, Scorpaena pepo*
- outside reserves, but not inside: *Aplodactylus arctidens, Callanthias japonicus, Chelidonichthys kumu, Chelidonichthys spinosus, Engraulis japonicus, Forsterygion maryannae, Gobiesox maeandricus, Hypoplectrodes huntii, Latridopsis ciliaris, Microcanthus strigatus, Odax pullus, Opistognathus punctatus, Opistognathus sp., Phoxinus sp., Pseudophycis barbata, Thyrsites atun, Trachurus japonicus*

[1] observations ordered by descending read abundance

[2] Taxon lists follow NCBI taxonomy (Federhen, 2012). Genera and species are known in Aotearoa / New Zealand (CD Roberts et al., 2019) unless indicated with an asterisk ("$^{*}$") or not a "fish" ("♠"), or freshwater taxa ("♣").

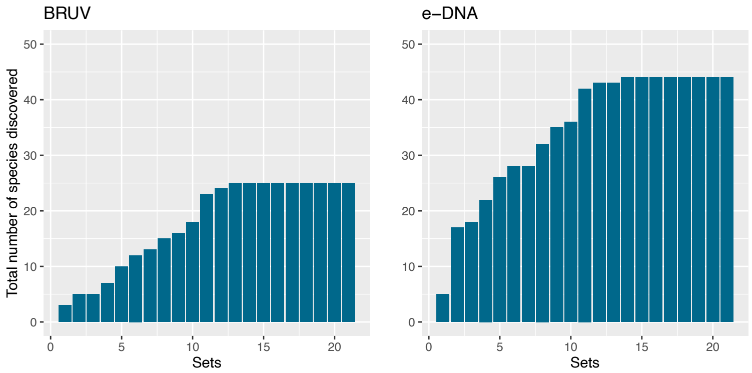

**Figure 5:** Species accumulation curves for BRUV and eDNA

- both inside and outside marine reserves: *Aldrichetta forsteri, Asterropteryx semipunctata, Bodianus unimaculatus, Bostrychus zonatus, Caesioperca lepidoptera, Caprodon schlegelii, Cheilodactylus variegatus, Cheilodactylus, zonatus, Coptodon zillii, Galaxias sp.* ('southern')*, Helcogramma striata, Helicolenus percoides, Lotella phycis, Lotella rhacina, Maurolicus muelleri, Meuschenia scaber, Nemadactylus macropterus, Notolabrus celidotus, Notolabrus cinctus, Opistognathus iyonis, Opistognathus liturus, Parapercis colias, Parapercis decemfasciata, Pseudolabrus miles, Scobinichthys granulatus*

#### Species accumulation curves

Species accumulation curves for BRUV and eDNA data are shown in Fig. 5.

#### DNA sequence alignments

Amplicon sequence variants affected by gaps in alignments for taxonomic assignments included, in part, those belonging to families Callanthiidae, Chaetodontidae, Cichlidae, Eleotridae, Gaidropsaridae, Gobiesocidae, Gobiidae, Kyphosidae, Lutjanidae, Monocentridae, Myctophidae, Opistognathidae, Percidae, Pinguipedidae, Serranidae, Sternoptychidae, and Tripterygiidae.

Amplicon sequence variants affected by imperfect query coverage included, in part, those belonging to families Anguillidae, Banjosidae, Bovichtidae, Callanthiidae, Carangidae, Chaetodontidae, Cheilodactylidae, Cichlidae, Cyprinidae, Eleotridae, Engraulidae, Gaidropsaridae, Galaxiidae, Gobiesocidae, Gobiidae, Hexanchidae, Kyphosidae, Lutjanidae, Merlucciidae, Monacanthidae, Monocentridae, Moridae, Mugilidae, Myctophidae, Opistognathidae, Percidae, Pinguipedidae, Salmonidae, Scombridae, Scorpaenidae, Sebastidae, Serranidae, Squalidae, Sternoptychidae, Triakidae, Triglidae, and Tripterygiidae.

Alignments for *Arctocephalus forsteri* (New Zealand fur seal), *Balaenoptera musculus* (blue whale), and *Tursiops truncatus* (bottlenose dolphin) are shown in Fig. 6 on the following page.

#### ANOSIM results

ANOSIM results are summarized in Table 3 on page 11.

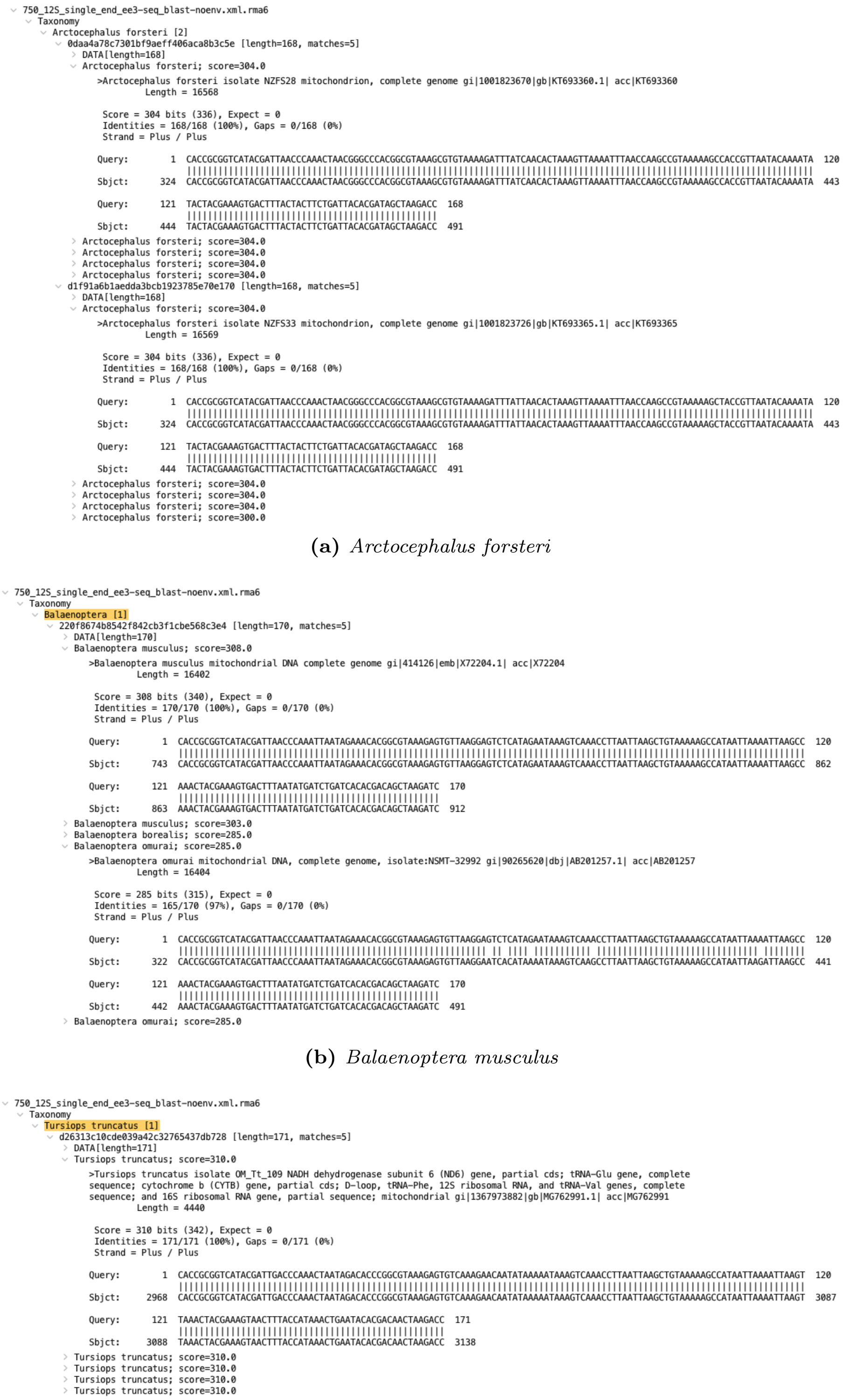

**(a)** *Arctocephalus forsteri*

**(b)** *Balaenoptera musculus*

**(c)** *Tursiops truncatus*

**Figure 6:** Alignment examples (a) for *Arctocephalus forsteri* (New Zealand fur seal), (b) *Balaenoptera musculus* (blue whale), and (c) *Tursiops truncatus* (bottlenose dolphin), as exported from Megan (Huson et al., 2016).

**Table 3:** Summary of ANOSIM (Clarke, 1993) results. Results are shown for Jaccard (Jaccard, 1912) distances of fish species observations between field work data sets, based on different grouping variables. Sets aggregated by grouping variables were used for permutation testing. Significant values derived from permutation testing within grouping variable (n = 9 999).

| Replication over | Tax. level | Location grouping | Obs. method | ANOSIM R | Significance |
|---|---|---|---|---|---|
| SET.ID | SPECIES | RESERVE.GROUP.LOCATION | eDNA | 0.113618524 | 0.1401 |
| | GENUS | | | 0.129317111 | 0.1066 |
| | FAMILY | | | 0.116758242 | 0.1349 |
| | ORDER | | | 0.090266876 | 0.1763 |
| | SPECIES | RESERVE.GROUP.INSIDE | | -0.075454545 | 0.8757 |
| | GENUS | | | -0.042090909 | 0.7155 |
| | FAMILY | | | -0.030363636 | 0.6528 |
| | ORDER | | | 0.003454545 | 0.4354 |
| | SPECIES | RESERVE.GROUP.LOCATION | BRUV | 0.383437991 | 0.0005 |
| | GENUS | | | 0.410518053 | 0.0003 |
| | FAMILY | | | 0.410518053 | 0.0004 |
| | ORDER | | | 0.289246468 | 0.0039 |
| | SPECIES | RESERVE.GROUP.INSIDE | | 0.106272727 | 0.0758 |
| | GENUS | | | 0.075818182 | 0.1296 |
| | FAMILY | | | 0.095727273 | 0.0855 |
| | ORDER | | | 0.064727273 | 0.1503 |

#### Indicator species analysis

Indicator species analysis saw *Bodianus unimaculatus* (red hogfish) non-randomly common at WJ CTRL+WJ MR (stat 0.725, $p$ 0.0282). On genus level, *Bodianus* (hogfish) appeared non-randomly at WJ CTRL+WJ MR (statistic 0.725, $p$ 0.03). At family level, Labridae (wrasses) appeared associated with FF CTRL+FF MR+WJ CTRL+WJ MR (stat 0.725, $p$ 0.0385). At order level Perciformes was common at FF MR+WJ CTRL+WJ MR (statistic 0.699, $p$ 0.0301).

#### Binomial regression

Regression analysis is summarized in Fig. 7 on the following page. Model coefficients are summarized in Fig. 8 on the next page.

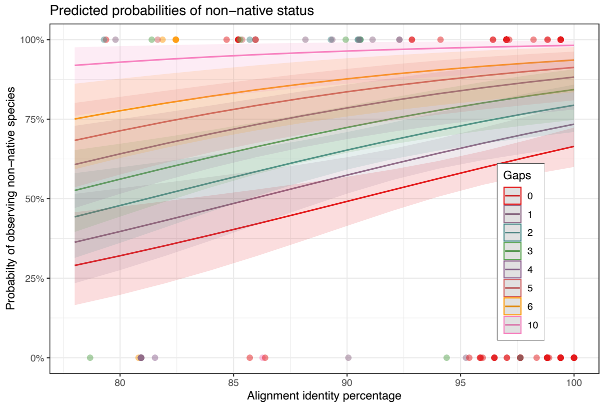

**Figure 7:** Summary of binomial regression. The response variable was non-native status of eDNA detected species (`TRUE`: eDNA derived species assignment not in CD Roberts et al. (2019), 19 species among 53 ASVs; `FALSE`: 25 species among 39 ASVs), predictor variables were the number of alignment gaps and alignment query coverage (%), the number of trials were defined using by numbers of locations at which each assigned species was seen. Graph created using R package *sjPlot* (2.8.0).

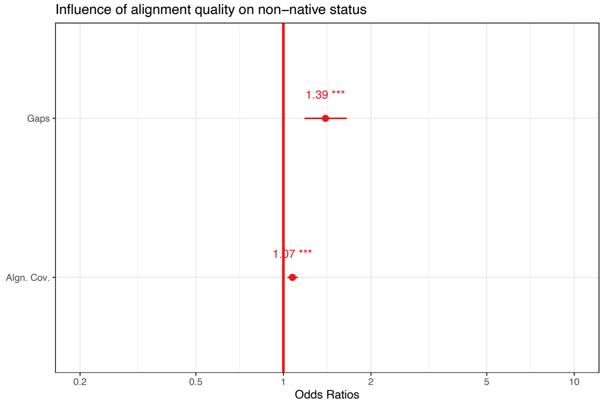

**Figure 8:** Relationship between non-native status (CD Roberts et al., 2019), gaps, and alignment coverage. The further the odds ratio from 1, the more pronounced the effect.

Bolyen, E., Rideout, J. R., Dillon, M. R., Bokulich, N. A., Abnet, C. C., Al-Ghalith, G. A., … Caporaso, J. G. (2019). Reproducible, interactive, scalable and extensible microbiome data science using QIIME 2. *Nature Biotechnology*, *37*(8), 852–857. doi:10/gf5292

Bylemans, J., Gleeson, D. M., Hardy, C. M., & Furlan, E. (2018). Toward an ecoregion scale evaluation of eDNA metabarcoding primers: A case study for the freshwater fish biodiversity of the murray-darling basin (australia). *Ecology and Evolution*, 1–16. doi:10/gfdrgt

Callahan, B., & Davis, N. M. (2021). *Decontam: Identify contaminants in marker-gene and metagenomics sequencing data*. R package version 1.12.0. Retrieved from https://github.com/benjjneb/decontam

Callahan, B. J., McMurdie, P. J., & Holmes, S. P. (2017). Exact sequence variants should replace operational taxonomic units in marker-gene data analysis. *The ISME Journal*, *11*(12), 113597. ISBN: 1751-7370 Publisher: Nature Publishing Group. doi:10/gcnwrw

Callahan, B. J., McMurdie, P. J., Rosen, M. J., Han, A. W., Johnson, A. J. A., & Holmes, S. P. (2016). DADA2: High-resolution sample inference from illumina amplicon data. *Nature Methods*, *13*(7), 581–583. ISBN: 1548-7105 (Electronic)$\backslash$r1548-7091 (Linking) _eprint: 15334406. doi:10/f84fxp

Camacho, C., Coulouris, G., Avagyan, V., Ma, N., Papadopoulos, J., Bealer, K., & Madden, T. L. (2009). BLAST+: Architecture and applications. *BMC Bioinformatics*, *10*(1), 421. ISBN: 1471210510. doi:10/cnjxgz

Chamberlain, S., Szoecs, E., Foster, Z., & Arendsee, Z. (2020). *Taxize: Taxonomic information from around the web*. R package version 0.9.99. Retrieved from https://CRAN.R-project.org/package=taxize

Clarke, K. R. (1993). Non-parametric multivariate analyses of changes in community structure. *Austral Ecology*, *18*(1), 117–143. doi:10.1111/j.1442-9993.1993.tb00438.x

De Cáceres, M., Legendre, P., & Moretti, M. (2010). Improving indicator species analysis by combining groups of sites. *Oikos*, *119*(10), 1674–1684. doi:10.1111/j.1600-0706.2010.18334.x

DiBattista, J. D., Coker, D. J., Sinclair-Taylor, T. H., Stat, M., Berumen, M. L., & Bunce, M. (2017). Assessing the utility of eDNA as a tool to survey reef-fish communities in the Red Sea. *Coral Reefs*, *36*(4), 1245–1252. doi:10.1007/s00338-017-1618-1

Dowle, M., & Srinivasan, A. (2021). *Data.table: Extension of ‘data.frame‘*. R package version 1.14.0. Retrieved from https://CRAN.R-project.org/package=data.table

Edgar, R. C., & Flyvbjerg, H. (2015). Error filtering, pair assembly and error correction for next-generation sequencing reads. *Bioinformatics*, *31*(21), 3476–3482. ISBN: 1367-4803. doi:10/gdq95z

Evans, N. T., Olds, B. P., Renshaw, M. A., Turner, C. R., Li, Y., Jerde, C. L., … Lodge, D. M. (2016). Quantification of mesocosm fish and amphibian species diversity via environmental DNA metabarcoding. *Molecular Ecology Resources*, *16*(1), 29–41. ISBN: 1755-098X. doi:10/gg3njr

Ewels, P., Magnusson, M., Lundin, S., & KÃ€ller, M. (2016). MultiQC: Summarize analysis results for multiple tools and samples in a single report. *Bioinformatics*, *32*(19), 3047–3048. doi:10/f3s996

Federhen, S. (2012). The NCBI taxonomy database. *Nucleic Acids Research*, *40*, D136–D143. doi:10/c452q3

Firke, S. (2021). *Janitor: Simple tools for examining and cleaning dirty data*. R package version 2.1.0. Retrieved from https://github.com/sfirke/janitor

Gohel, D. (2021a). *Flextable: Functions for tabular reporting*. R package version 0.6.7. Retrieved from https://CRAN.R-project.org/package=flextable

Gohel, D. (2021b). *Officer: Manipulation of microsoft word and powerpoint documents.* R package version 0.3.19. Retrieved from https://CRAN.R-project.org/package=officer

Grange, K. R. (1985). *The intertidal ecology of the soft shores of freshwater basin, milford sound. report prepared by the new zealand oceanographic institute for the department of lands and survey.* New Zealand Oceanographic Institute. Wellington, New Zealand.

Hester, J. (2020). *Primertree: Visually assessing the specificity and informativeness of primer pairs.* R package version 1.0.5. Retrieved from https://CRAN.R-project.org/package=primerTree

Huson, D. H., Beier, S., Flade, I., Gï¿œrska, A., El-Hadidi, M., Mitra, S., ... Tappu, R. (2016). MEGAN community edition - interactive exploration and analysis of large-scale microbiome sequencing data. *PLOS Computational Biology*, *12*(6), e1004957. doi:10/gfzpks

Inglis, G., Post-border Directorate, & MAF Biosecurity New Zealand. (2008). *Milford sound: First baseline survey for non-indigenous marine species (research project ZBS2005/19).* OCLC: 702296120. Wellington, N.Z.: MAF Biosecurity New Zealand. Retrieved August 5, 2021, from http://www.biosecurity.govt.nz/files/pests/salt-freshwater/milford-2008-resurvey-report.pdf

Ivanova, N. V., Zemlak, T. S., Hanner, R. H., & Hebert, P. D. N. (2007). Universal primer cocktails for fish DNA barcoding. *Molecular Ecology Notes*, *7*(4), 544–548. doi:10.1111/j.1471-8286.2007.01748.x

Jaccard, P. (1912). The distribution of the flora in the alpine zone. *New Phytologist*, *11*(2), 37–50. doi:10/fvhsjd

Jeunen, G.-J., Urban, L., Lewis, R., Knapp, M., Lamare, M., Rayment, W., ... Gemmell, N. (2020). Marine environmental DNA (eDNA) for biodiversity assessments: A one-to-one comparison between eDNA and baited remote underwater video (BRUV) surveys. doi:10.22541/au.160278512.26241559/v1

Kassambara, A. (2020). *Ggpubr: Ggplot2 based publication ready plots.* R package version 0.4.0. Retrieved from https://rpkgs.datanovia.com/ggpubr/

Kelly, R. P., Port, J. A., Yamahara, K. M., & Crowder, L. B. (2014). Using Environmental DNA to Census Marine Fishes in a Large Mesocosm. *PLOS ONE*, *9*(1), e86175. doi:10.1371/journal.pone.0086175

Kitano, T., Umetsu, K., Tian, W., & Osawa, M. (2007). Two universal primer sets for species identification among vertebrates. *International Journal of Legal Medicine*, *121*(5), 423–427. doi:10.1007/s00414-006-0113-y

Kocher, T. D., Thomas, W. K., Meyer, A., Edwards, S. V., Pï¿œï¿œbo, S., Villablanca, F. X., & Wilson, A. C. (1989). Dynamics of mitochondrial DNA evolution in animals: Amplification and sequencing with conserved primers. *Proceedings of the National Academy of Sciences*, *86*(16), 6196–6200. Publisher: National Academy of Sciences Section: Research Article. doi:10/c2hk4r

Lüdecke, D. (2021). *Sjplot: Data visualization for statistics in social science.* R package version 2.8.9. Retrieved from https://strengejacke.github.io/sjPlot/

Majaneva, M., Diserud, O. H., Eagle, S. H. C., Bostrï¿œm, E., Hajibabaei, M., & Ekrem, T. (2018). Environmental DNA filtration techniques affect recovered biodiversity. *Scientific Reports*, *8*(1), 4682. ISBN: 4159801823052. doi:10/gc7tf2

Marques, V., Milhau, T., Albouy, C., Dejean, T., Manel, S., Mouillot, D., & Juhel, J.-B. (2021). GAPeDNA: Assessing and mapping global species gaps in genetic databases for eDNA metabarcoding. *Diversity and Distributions*. Place: Hoboken Publisher: Wiley WOS:000629615300001. doi:10.1111/ddi.13142

Martin, M. (2011). Cutadapt removes adapter sequences from high-throughput sequencing reads. *EMBnet.journal*, *17*(1), 10. _eprint: ISSN 2226-6089. doi:10.14806/ej.17.1.200

McInnes, J. C., Jarman, S. N., Lea, M.-A., Raymond, B., Deagle, B. E., Phillips, R. A., … Alderman, R. (2017). DNA Metabarcoding as a Marine Conservation and Management Tool: A Circumpolar Examination of Fishery Discards in the Diet of Threatened Albatrosses. *Frontiers in Marine Science*, *4*, 277. doi:10.3389/fmars.2017.00277

McMurdie, P. J., Holmes, S., with contributions from Gregory Jordan, & Chamberlain, S. (2021). *Phyloseq: Handling and analysis of high-throughput microbiome census data*. R package version 1.36.0. Retrieved from http://dx.plos.org/10.1371/journal.pone.0061217

Miya, M., Sato, Y., Fukunaga, T., Sado, T., Poulsen, J. Y., Sato, K., … Iwasaki, W. (2015). MiFish, a set of universal PCR primers for metabarcoding environmental DNA from fishes: Detection of more than 230 subtropical marine species. *Royal Society Open Science*, *2*(7), 150088. ISBN: 2054-5703. doi:10/gmcj95

Mladenov, P. V. (2001). New zealand fiords: Researching, managing, and conserving a unique ecosystem. *New Zealand Journal of Marine and Freshwater Research*, *35*(4), 653–661. doi:10.1080/00288330.2001.9517032

Nicholson, J. M., Mordaunt, M., Lopez, P., Uppala, A., Rosati, D., Rodrigues, N. P., … Rife, S. C. (2021). *Scite: A smart citation index that displays the context of citations and classifies their intent using deep learning*. Cold Spring Harbor Laboratory. doi:10.1101/2021.03.15.435418

Oksanen, J., Blanchet, F. G., Kindt, R., Legendre, P., Minchin, P. R., O'Hara, R. B., … Wagner, H. (2015). Vegan: Community ecology package. CRAN.

Ooms, J. (2021a). *Curl: A modern and flexible web client for r*. R package version 4.3.2. Retrieved from https://CRAN.R-project.org/package=curl

Ooms, J. (2021b). *Magick: Advanced graphics and image-processing in r*. R package version 2.7.3. Retrieved from https://CRAN.R-project.org/package=magick

Palumbi, S. R. (1996). Nucleic Acids II: The Polymerase Chain Reaction. In *Molecular Systematics* (Second, Vol. 2). Sunderland, Massachusetts: Sinauer Associates.

Pebesma, E. (2021). *Sf: Simple features for r*. R package version 1.0-2. Retrieved from https://CRAN.R-project.org/package=sf

Pebesma, E., & Bivand, R. (2021). *Sp: Classes and methods for spatial data*. R package version 1.4-5. Retrieved from https://CRAN.R-project.org/package=sp

Provoost, P., & Bosch, S. (2021). *Robis: Ocean biodiversity information system (obis) client*. R package version 2.6.1. Retrieved from https://github.com/iobis/robis

R Core Development Team. (2019). R: A language and environment for statistical computing. R Foundation for Statistical Computing. Retrieved from http://www.r-project.org/.

Roberts, C. [CD], Stewart, A., Struthers, C., Barker, J., & Kortet, S. (2019). *Checklist of the fishes of new zealand* (No. July). Museum of New Zealand Te Papa Tongarewa. Series: Version 1.1 July 2019. Retrieved from https://collections.tepapa.govt.nz/document/10564

Roberts, C. [Clive] (Ed.). (2005). *Regional diversity and biogeography of coastal fishes on the west coast south island of new zealand*. Wellington, N.Z: Dept. of Conservation, 250.

Rosen, M. J., Callahan, B. J., Fisher, D. S., & Holmes, S. P. (2012). Denoising PCR-amplified metagenome data. *BMC Bioinformatics*, *13*(1). ISBN: 1471-2105 (Electronic)$\backslash$r1471-2105 (Linking). doi:10/gb8vq2

Schauberger, P., & Walker, A. (2021). *Openxlsx: Read, write and edit xlsx files*. R package version 4.2.4. Retrieved from https://CRAN.R-project.org/package=openxlsx

Schnell, I. B., Bohmann, K., & Gilbert, M. T. P. (2015). Tag jumps illuminated - reducing sequence-to-sample misidentifications in metabarcoding studies. *Molecular Ecology Resources*, *15*(6), 1289–1303. doi:10.1111/1755-0998.12402

Shaw, J. L. A., Clarke, L. J., Wedderburn, S. D., Barnes, T. C., Weyrich, L. S., & Cooper, A. (2016). Comparison of environmental DNA metabarcoding and conventional fish survey methods in a river system. *Biological Conservation*, *197*, 131–138. doi:10.1016/j.biocon.2016.03.010

Sherrill-Mix, S. (2021). *Taxonomizr: Functions to work with ncbi accessions and taxonomy*. R package version 0.8.0. Retrieved from https://CRAN.R-project.org/package=taxonomizr

Taberlet, P., Bonin, A., Zinger, L., & Coissac, E. (2018). *Environmental DNA: For biodiversity research and monitoring* (1st ed.). New York, NY: Oxford University Press.

Thomsen, P. F., Kielgast, J., Iversen, L. L., Møller, P. R., Rasmussen, M., & Willerslev, E. (2012). Detection of a Diverse Marine Fish Fauna Using Environmental DNA from Seawater Samples. *PLOS ONE*, *7*(8), e41732. doi:10.1371/journal.pone.0041732

Valentini, A., Taberlet, P., Miaud, C., Civade, R., Herder, J., Thomsen, P. F., … Dejean, T. (2016). Next-generation monitoring of aquatic biodiversity using environmental DNA metabarcoding. *Molecular Ecology*, *25*(4), 929–942. doi:10.1111/mec.13428

Ward, R. D., Zemlak, T. S., Innes, B. H., Last, P. R., & Hebert, P. D. (2005). DNA barcoding Australia's fish species. *Philosophical Transactions of the Royal Society B: Biological Sciences*, *360*(1462), 1847–1857. doi:10/c6vm58

Wickham, H. (2020). *Reshape2: Flexibly reshape data: A reboot of the reshape package*. R package version 1.4.4. Retrieved from https://github.com/hadley/reshape

Wickham, H. (2021a). *Tidyr: Tidy messy data*. R package version 1.1.3. Retrieved from https://CRAN.R-project.org/package=tidyr

Wickham, H. (2021b). *Tidyverse: Easily install and load the tidyverse*. R package version 1.3.1. Retrieved from https://CRAN.R-project.org/package=tidyverse

Wickham, H., & Bryan, J. (2019). *Readxl: Read excel files*. R package version 1.3.1. Retrieved from https://CRAN.R-project.org/package=readxl

Wickham, H., François, R., Henry, L., & Müller, K. (2021). *Dplyr: A grammar of data manipulation*. R package version 1.0.7. Retrieved from https://CRAN.R-project.org/package=dplyr

Wing, S. R., & Jack, L. (2013). Marine reserve networks conserve biodiversity by stabilizing communities and maintaining food web structure. *Ecosphere*, *4*(11), art135. doi:10.1890/ES13-00257.1

Xie, Y. (2021a). *Knitr: A general-purpose package for dynamic report generation in r*. R package version 1.33. Retrieved from https://yihui.org/knitr/

Xie, Y. (2021b). *Xaringan: Presentation ninja*. R package version 0.22. Retrieved from https://github.com/yihui/xaringan

### A Taxonomic observations

**Table 4:** Details on taxonomic observations across data sources. Taxonomic hierarchies conform with NCBI taxonomy, where available, and are sorted alphabetically – the resulting species order is identical to Fig. 2 of *main text*. Taxa not listed as New Zealand species in (CD Roberts et al., 2019) are highlighted with Asterisk (*). Trivial names are indicated, where available from NCBI.

| Class | Order | Family | Genus | Species | Common name | BRUV | eDNA | OBIS | Literature |
|---|---|---|---|---|---|---|---|---|---|
| Actinopteri | Anguilliformes | Anguillidae | *Anguilla* | *Anguilla australis* | Australian shortfin eel | | 1 | | |
| | | Congridae | *Conger* | *Conger verreauxi* | conger eel | | | | TRUE |
| | Atheriniformes | Atherinidae | *Atherinomorus* | *Atherinomorus lacunosus* | hardyhead silverside | | | | TRUE |
| | Blenniiformes | Gobiesocidae | *Gobiesox** | *Gobiesox maeandricus** | northern clingfish | | 1 | | |
| | | Tripterygiidae | *Bellapiscis* | *Bellapiscis lesleyae* | mottled twister | | | | TRUE |
| | | | | *Bellapiscis medius* | twister | | | | TRUE |
| | | | *Cryptichthys* | *Cryptichthys jojettae* | | | | | TRUE |
| | | | *Forsterygion* | *Forsterygion capito* | spotted robust triplefin | | | | TRUE |
| | | | | *Forsterygion flavonigrum* | yellow-and-black triplefin | | | TRUE | TRUE |
| | | | | *Forsterygion lapillum* | common triplefin | | | | TRUE |
| | | | | *Forsterygion malcolmi* | | | | | TRUE |
| | | | | *Forsterygion maryannae* | | 4 | | | TRUE |
| | | | | *Forsterygion varium* | striped triplefin | | | | TRUE |
| | | | *Helcogramma** | *Helcogramma striata** | | | 6 | | |
| | | | *Karalepis* | *Karalepis stewarti* | | | | | TRUE |
| | | | *Notoclinops* | *Notoclinops caerulepunctus* | | | | | TRUE |
| | | | | *Notoclinops segmentatus* | | | | | TRUE |
| | | | *Notoclinus* | *Notoclinus compressus* | | | | TRUE | |
| | | | | *Notoclinus fenestratus* | | | | | TRUE |
| | | | *Ruanoho* | *Ruanoho decemdigitatus* | | | | | TRUE |
| | | | | *Ruanoho whero* | spectacled triplefin | | | | TRUE |
| | Carangiformes | Carangidae | *Trachurus* | *Trachurus japonicus* | Japanese jack mackerel | | 1 | | |
| | Centrarchiformes | Aplodactylidae | *Aplodactylus* | *Aplodactylus arctidens* | | 2 | | | TRUE |
| | | Cheilodactylidae | *Cheilodactylus* | *Cheilodactylus variegatus* | | | 7 | | |
| | | | | *Cheilodactylus zonatus* | blackbarred morwong | | 7 | | |
| | | | *Nemadactylus* | *Nemadactylus macropterus* | tarakihi | 14 | | TRUE | TRUE |
| | | Kyphosidae | *Microcanthus** | *Microcanthus strigatus** | stripey | | 1 | | |

| Class (cont.) | Order (cont.) | Family (cont.) | Genus (cont.) | Species (cont.) | Common name (cont.) | BRUV (cont.) | eDNA (cont.) | OBIS (cont.) | Literature (cont.) |
|---|---|---|---|---|---|---|---|---|---|
| | | | *Scorpis* | *Scorpis lineolata* | silver sweep | | | | TRUE |
| | | Latridae | *Latridopsis* | *Latridopsis ciliaris* | blue moki | 1 | | TRUE | TRUE |
| | | | | *Latridopsis forsteri* | bastard trumpeter | | | | TRUE |
| | | | *Latris* | *Latris lineata* | striped trumpeter | | | TRUE | TRUE |
| | | | *Mendosoma* | *Mendosoma lineatum* | | | | | TRUE |
| | Chaetodontiformes | Chaetodontidae | *Chaetodon* | *Chaetodon zanzibarensis* | | | 1 | | |
| | Cichliformes | Cichlidae | *Benitochromis** | *Benitochromis finleyi** | | | 1 | | |
| | | | *Coptodon** | *Coptodon zillii** | redbelly tilapia | | 2 | | |
| | Clupeiformes | Engraulidae | *Engraulis** | *Engraulis japonicus** | Japanese anchovy | | 1 | | |
| | Cypriniformes | Cyprinidae | *Phoxinus** | *Phoxinus sp.** | | | 1 | | |
| | Gadiformes | Gaidropsaridae | *Gaidropsarus* | *Gaidropsarus argentatus* | Arctic rockling | | 1 | | |
| | | | | *Gaidropsarus novaezelandi* | | | | | TRUE |
| | | Merlucciidae | *Macruronus* | *Macruronus novaezelandiae* | blue grenadier | | 1 | | |
| | | Moridae | *Lotella* | *Lotella phycis* | | | 2 | | |
| | | | | *Lotella rhacina* | rock cod | 2 | | | TRUE |
| | | | *Pseudophycis* | *Pseudophycis barbata* | southern bastard codling | 1 | | TRUE | TRUE |
| | Galaxiiformes | Galaxiidae | *Galaxias* | *Galaxias argenteus* | | | | TRUE | |
| | | | | *Galaxias sp.* | | | 4 | | |
| | Gobiesociformes | Gobiesocidae | *Modicus* | *Modicus minimus* | | | | | TRUE |
| | | | | *Modicus tangaroa* | | | | | TRUE |
| | Gobiiformes | Eleotridae | *Bostrychus** | *Bostrychus zonatus** | barred gudgeon | | 8 | | |
| | | Gobiidae | *Asterropteryx** | *Asterropteryx semipunctata** | starry goby | | 5 | | |
| | | | *Gobiopsis* | *Gobiopsis atrata* | | | | | TRUE |
| | | Thalasseleotrididae | *Thalasseleotris* | *Thalasseleotris iota* | | | | TRUE | |
| | Labriformes | Labridae | *Bodianus* | *Bodianus unimaculatus* | red pigfish | 5 | | | |
| | | | *Notolabrus* | *Notolabrus celidotus* | New Zealand spotty | 5 | | TRUE | TRUE |
| | | | | *Notolabrus cinctus* | | 4 | | TRUE | TRUE |
| | | | | *Notolabrus fucicola* | yellow-saddled wrasse | 1 | | TRUE | TRUE |
| | | | *Pseudolabrus* | *Pseudolabrus miles* | | 16 | | TRUE | TRUE |
| | | Odacidae | *Odax* | *Odax pullus* | greenbone | 1 | | | TRUE |

| Class (cont.) | Order (cont.) | Family (cont.) | Genus (cont.) | Species (cont.) | Common name (cont.) | BRUV (cont.) | eDNA (cont.) | OBIS (cont.) | Literature (cont.) |
|---|---|---|---|---|---|---|---|---|---|
| | Lutjaniformes | Lutjanidae | *Lutjanus* | *Lutjanus sanguineus* | humphead snapper | | 1 | | |
| | Mugiliformes | Mugilidae | *Aldrichetta* | *Aldrichetta forsteri* | yellow-eye mullet | | 2 | | TRUE |
| | Myctophiformes | Myctophidae | *Gymnoscopelus** | *Gymnoscopelus nicholsi** | | | 1 | | |
| | | | *Hygophum* | *Hygophum hygomii* | | | 1 | | |
| | Ophidiiformes | Bythitidae | *Fiordichthys* | *Fiordichthys slartibartfasti* | | | | | TRUE |
| | Osmeriformes | Retropinnidae | *Retropinna* | *Retropinna retropinna* | cucumberfish | | | | TRUE |
| | Ovalentaria | Plesiopidae | *Acanthoclinus* | *Acanthoclinus fuscus* | | | | | TRUE |
| | | | | *Acanthoclinus littoreus* | | | | | TRUE |
| | | | | *Acanthoclinus marilynae* | | | | | TRUE |
| | | | | *Acanthoclinus matti* | | | | TRUE | TRUE |
| | | | | *Acanthoclinus rua* | | | | | TRUE |
| | Pempheriformes | Banjosidae | *Banjos** | *Banjos banjos** | | | 6 | | |
| | | Percophidae | *Hemerocoetes* | *Hemerocoetes monopterygius* | | | | TRUE | |
| | | Polyprionidae | *Polyprion* | *Polyprion oxygeneios* | | | | | TRUE |
| | Perciformes | Bovichtidae | *Bovichtus** | *Bovichtus diacanthus** | | | 1 | | |
| | | | | *Bovichtus variegatus** | thornfish | | | | TRUE |
| | | Callanthiidae | *Callanthias* | *Callanthias allporti* | | | | | TRUE |
| | | | | *Callanthias japonicus* | | | 1 | | |
| | | Nototheniidae | *Notothenia* | *Notothenia angustata* | Maori chief | | | TRUE | |
| | | Percidae | *Sander** | *Sander lucioperca** | pikeperch | | 1 | | |
| | | Scorpaenidae | *Scorpaena* | *Scorpaena cardinalis* | red rock cod | 1 | | | |
| | | | | *Scorpaena papillosa* | | | | TRUE | TRUE |
| | | | | *Scorpaena pepo* | pumpkin scorpionfish | | 1 | | |
| | | Sebastidae | *Helicolenus* | *Helicolenus hilgendorfi* | | | 1 | | |
| | | | | *Helicolenus percoides* | | 6 | | | TRUE |
| | | Serranidae | *Caesioperca* | *Caesioperca lepidoptera* | | 8 | | TRUE | TRUE |
| | | | *Caprodon** | *Caprodon schlegelii** | sunrise perch | | 27 | | |
| | | | *Hypoplectrodes* | *Hypoplectrodes huntii* | | 2 | | | TRUE |
| | | | *Lepidoperca* | *Lepidoperca tasmanica* | | | | TRUE | TRUE |
| | | Triglidae | *Chelidonichthys* | *Chelidonichthys kumu* | bluefin gurnard | 1 | | | |

| Class (cont.) | Order (cont.) | Family (cont.) | Genus (cont.) | Species (cont.) | Common name (cont.) | BRUV (cont.) | eDNA (cont.) | OBIS (cont.) | Literature (cont.) |
|---|---|---|---|---|---|---|---|---|---|
| | | | | *Chelidonichthys spinosus* | red gurnard | | 1 | | |
| | Pleuronectiformes | Rhombosoleidae | *Peltorhamphus* | *Peltorhamphus latus* | speckled sole | | | TRUE | |
| | | | *Rhombosolea* | *Rhombosolea plebeia* | New Zealand flounder | | | | TRUE |
| | Salmoniformes | Salmonidae | *Oncorhynchus* | *Oncorhynchus mykiss* | rainbow trout | | 1 | | |
| | Scombriformes | Gempylidae | *Thyrsites* | *Thyrsites atun* | snoek | 1 | | | TRUE |
| | | Scombridae | *Katsuwonus* | *Katsuwonus pelamis* | skipjack tuna | | 1 | | |
| | | | *Scomber* | *Scomber japonicus* | chub mackerel | | 1 | | |
| | Stomiiformes | Sternoptychidae | *Maurolicus* | *Maurolicus muelleri* | pearlsides | | 9 | | |
| | Tetraodontiformes | Monacanthidae | *Meuschenia* | *Meuschenia scaber* | velvet leatherjacket | 6 | | TRUE | TRUE |
| | | | *Scobinichthys** | *Scobinichthys granulatus** | rough leatherjacket | | 11 | | |
| | Trachichthyiformes | Monocentridae | *Monocentris* | *Monocentris japonicus* | | | 2 | | |
| | | Trachichthyidae | *Paratrachichthys* | *Paratrachichthys trailli* | sandpaper fish | | | | TRUE |
| | undefined | Opistognathidae | *Opistognathus** | *Opistognathus iyonis** | | | 3 | | |
| | | | | *Opistognathus liturus** | seto-amadai | | 2 | | |
| | | | | *Opistognathus punctatus** | finespotted jawfish | | 1 | | |
| | | | | *Opistognathus sp.** | | | 1 | | |
| | Uranoscopiformes | Pinguipedidae | *Parapercis* | *Parapercis colias* | New Zealand blue cod | 21 | | TRUE | TRUE |
| | | | | *Parapercis decemfasciata* | | | 10 | | |
| | | | | *Parapercis gilliesii* | yellow weaver | | | | TRUE |
| Chondrichthyes | Carcharhiniformes | Carcharhinidae | *Prionace* | *Prionace glauca* | blue shark | | | TRUE | |
| | | Scyliorhinidae | *Cephaloscyllium* | *Cephaloscyllium isabellum* | | 4 | | | TRUE |
| | | Triakidae | *Galeorhinus* | *Galeorhinus galeus* | tope shark | 2 | | | |
| | | | *Mustelus* | *Mustelus lenticulatus* | spotted estuary smooth-hound | 1 | | | |
| | | | | *Mustelus manazo* | starspotted smooth-hound | | 1 | | |
| | Hexanchiformes | Hexanchidae | *Notorynchus* | *Notorynchus cepedianus* | broadnose sevengill shark | 2 | 1 | | |
| | Lamniformes | Alopiidae | *Carcharodon* | *Carcharodon carcharias* | great white shark | | | TRUE | |
| | | | *Isurus* | *Isurus oxyrinchus* | shortfin mako shark | | | TRUE | |
| | Squaliformes | Squalidae | *Squalus* | *Squalus acanthias* | spiny dogfish | 2 | | TRUE | TRUE |
| | | | | *Squalus suckleyi* | Puget Sound dogfish | | 3 | | |
