## Supplementary material for "Environmental DNA analysis needs local reference data to inform taxonomy-based conservation policy – A case study from Aotearoa / New Zealand": Table 1

| Class | Order | Family | Genus | Species | Common name | Algn. covrg. | Algn. gaps |
|---|---|---|---|---|---|---|---|
| Actinopteri | Anguilliformes | Anguillidae | *Anguilla* | *Anguilla australis* | Australian shortfin eel | 100% | 0 |
| | | Congridae | *Conger* | *Conger verreauxi* | conger eel | | |
| | Atheriniformes | Atherinidae | *Atherinomorus* | *Atherinomorus lacunosus* | hardyhead silverside | | |
| | Blenniiformes | Gobiesocidae | *Gobiesox** | *Gobiesox maeandricus** | northern clingfish | 79.30% | 2 |
| | | Tripterygiidae | *Bellapiscis* | *Bellapiscis lesleyae* | mottled twister | | |
| | | | | *Bellapiscis medius* | twister | | |
| | | | *Cryptichthys* | *Cryptichthys jojettae* | | | |
| | | | *Forsterygion* | *Forsterygion capito* | spotted robust triplefin | | |
| | | | | *Forsterygion flavonigrum* | yellow-and-black triplefin | | |
| | | | | *Forsterygion lapillum* | common triplefin | | |
| | | | | *Forsterygion malcolmi* | | | |
| | | | | *Forsterygion maryannae* | | | |
| | | | | *Forsterygion varium* | striped triplefin | | |
| | | | *Helcogramma** | *Helcogramma striata** | | 88.2-92.3% | 1 |
| | | | *Karalepis* | *Karalepis stewarti* | | | |
| | | | *Notoclinops* | *Notoclinops caerulepunctus* | | | |
| | | | | *Notoclinops segmentatus* | | | |
| | | | *Notoclinus* | *Notoclinus compressus* | | | |
| | | | | *Notoclinus fenestratus* | | | |
| | | | *Ruanoho* | *Ruanoho decemdigitatus* | | | |
| | | | | *Ruanoho whero* | spectacled triplefin | | |
| | Carangiformes | Carangidae | *Trachurus* | *Trachurus japonicus* | Japanese jack mackerel | 99.40% | 0 |
| | Centrarchiformes | Aplodactylidae | *Aplodactylus* | *Aplodactylus arctidens* | | | |
| | | Cheilodactylidae | *Cheilodactylus* | *Cheilodactylus variegatus* | | 97.6-98.8% | 0 |
| | | | | *Cheilodactylus zonatus* | blackbarred morwong | 97-97.6% | 0 |
| | | | *Nemadactylus* | *Nemadactylus macropterus* | tarakihi | | |
| | | Kyphosidae | *Microcanthus** | *Microcanthus strigatus** | stripey | 85.30% | 3 |
| | | | *Scorpis* | *Scorpis lineolata* | silver sweep | | |
| | | Latridae | *Latridopsis* | *Latridopsis ciliaris* | blue moki | | |
| | | | | *Latridopsis forsteri* | bastard trumpeter | | |
| | | | *Latris* | *Latris lineata* | striped trumpeter | | |
| | | | *Mendosoma* | *Mendosoma lineatum* | | | |
| | Chaetodontiformes | Chaetodontidae | *Chaetodon* | *Chaetodon zanzibarensis* | | 81.50% | 4 |
| | Cichliformes | Cichlidae | *Benitochromis** | *Benitochromis finleyi** | | 79.40% | 0 |
| | | | *Coptodon** | *Coptodon zillii** | redbelly tilapia | 90.60% | 3 |
| | Clupeiformes | Engraulidae | *Engraulis** | *Engraulis japonicus** | Japanese anchovy | 98.80% | 0 |
| | Cypriniformes | Cyprinidae | *Phoxinus** | *Phoxinus sp.** | | 97.10% | 0 |
| | Gadiformes | Gaidropsaridae | *Gaidropsarus* | *Gaidropsarus argentatus* | Arctic rockling | 90.10% | 1 |
| | | | | *Gaidropsarus novaezelandi* | | | |
| | | Merlucciidae | *Macruronus* | *Macruronus novaezelandiae* | blue grenadier | 100% | 0 |
| | | Moridae | *Lotella* | *Lotella phycis* | | 95.90% | 0 |
| | | | | *Lotella rhacina* | rock cod | | |
| | | | *Pseudophycis* | *Pseudophycis barbata* | southern bastard codling | | |

| Class (cont.) | Order (cont.) | Family (cont.) | Genus (cont.) | Species (cont.) | Common name (cont.) | Algn. covrg. (cont.) | Algn. gaps (cont.) |
|---|---|---|---|---|---|---|---|
| | Galaxiiformes | Galaxiidae | *Galaxias* | *Galaxias argenteus* | | | |
| | | | | *Galaxias sp.* | | 96.50% | 0 |
| | Gobiesociformes | Gobiesocidae | *Modicus* | *Modicus minimus* | | | |
| | | | | *Modicus tangaroa* | | | |
| | Gobiiformes | Eleotridae | *Bostrychus** | *Bostrychus zonatus** | barred gudgeon | 85.4-86% | 5 |
| | | Gobiidae | *Asterropteryx** | *Asterropteryx semipunctata** | starry goby | 81.9-82.5% | 6 |
| | | | *Gobiopsis* | *Gobiopsis atrata* | | | |
| | | Thalasseleotrididae | *Thalasseleotris* | *Thalasseleotris iota* | | | |
| | Labriformes | Labridae | *Bodianus* | *Bodianus unimaculatus* | red pigfish | | |
| | | | *Notolabrus* | *Notolabrus celidotus* | New Zealand spotty | | |
| | | | | *Notolabrus cinctus* | | | |
| | | | | *Notolabrus fucicola* | yellow-saddled wrasse | | |
| | | | *Pseudolabrus* | *Pseudolabrus miles* | | | |
| | | Odacidae | *Odax* | *Odax pullus* | greenbone | | |
| | Lutjaniformes | Lutjanidae | *Lutjanus* | *Lutjanus sanguineus* | humphead snapper | 80.80% | 6 |
| | Mugiliformes | Mugilidae | *Aldrichetta* | *Aldrichetta forsteri* | yellow-eye mullet | 96-100% | 0 |
| | Myctophiformes | Myctophidae | *Gymnoscopelus** | *Gymnoscopelus nicholsi** | | 79.80% | 1 |
| | | | *Hygophum* | *Hygophum hygomii* | | 78.70% | 3 |
| | Ophidiiformes | Bythitidae | *Fiordichthys* | *Fiordichthys slartibartfasti* | | | |
| | Osmeriformes | Retropinnidae | *Retropinna* | *Retropinna retropinna* | cucumberfish | | |
| | Ovalentaria | Plesiopidae | *Acanthoclinus* | *Acanthoclinus fuscus* | | | |
| | | | | *Acanthoclinus littoreus* | | | |
| | | | | *Acanthoclinus marilynae* | | | |
| | | | | *Acanthoclinus matti* | | | |
| | | | | *Acanthoclinus rua* | | | |
| | Pempheriformes | Banjosidae | *Banjos** | *Banjos banjos** | | 84.7-85.2% | 0 |
| | | Percophidae | *Hemerocoetes* | *Hemerocoetes monopterygius* | | | |
| | | Polyprionidae | *Polyprion* | *Polyprion oxygeneios* | | | |
| | Perciformes | Bovichtidae | *Bovichtus** | *Bovichtus diacanthus** | | 94.10% | 0 |
| | | | | *Bovichtus variegatus** | thornfish | | |
| | | Callanthiidae | *Callanthias* | *Callanthias allporti* | | | |
| | | | | *Callanthias japonicus* | | 95.20% | 1 |
| | | Nototheniidae | *Notothenia* | *Notothenia angustata* | Maori chief | | |
| | | Percidae | *Sander** | *Sander lucioperca** | pikeperch | 81.40% | 3 |
| | | Scorpaenidae | *Scorpaena* | *Scorpaena cardinalis* | red rock cod | | |
| | | | | *Scorpaena papillosa* | | | |
| | | | | *Scorpaena pepo* | pumpkin scorpionfish | 85.70% | 0 |
| | | Sebastidae | *Helicolenus* | *Helicolenus hilgendorfi* | | 95.40% | 0 |
| | | | | *Helicolenus percoides* | | | |
| | | Serranidae | *Caesioperca* | *Caesioperca lepidoptera* | | | |
| | | | *Caprodon** | *Caprodon schlegelii** | sunrise perch | 90.6-98.2% | 0-2 |
| | | | *Hypoplectrodes* | *Hypoplectrodes huntii* | | | |
| | | | *Lepidoperca* | *Lepidoperca tasmanica* | | | |

| Class (cont.) | Order (cont.) | Family (cont.) | Genus (cont.) | Species (cont.) | Common name (cont.) | Algn. covrg. (cont.) | Algn. gaps (cont.) |
|---|---|---|---|---|---|---|---|
| | | Triglidae | *Chelidonichthys* | *Chelidonichthys kumu* | bluefin gurnard | | |
| | | | | *Chelidonichthys spinosus* | red gurnard | 99.40% | 0 |
| | Pleuronectiformes | Rhombosoleidae | *Peltorhamphus* | *Peltorhamphus latus* | speckled sole | | |
| | | | *Rhombosolea* | *Rhombosolea plebeia* | New Zealand flounder | | |
| | Salmoniformes | Salmonidae | *Oncorhynchus* | *Oncorhynchus mykiss* | rainbow trout | 100% | 0 |
| | Scombriformes | Gempylidae | *Thyrsites* | *Thyrsites atun* | snoek | | |
| | | Scombridae | *Katsuwonus* | *Katsuwonus pelamis* | skipjack tuna | 95.90% | 0 |
| | | | *Scomber* | *Scomber japonicus* | chub mackerel | 100% | 0 |
| | Stomiiformes | Sternoptychidae | *Maurolicus* | *Maurolicus muelleri* | pearlsides | 86.3-99.4% | 0-10 |
| | Tetraodontiformes | Monacanthidae | *Meuschenia* | *Meuschenia scaber* | velvet leatherjacket | | |
| | | | *Scobinichthys** | *Scobinichthys granulatus** | rough leatherjacket | 98.8-99.4% | 0 |
| | Trachichthyiformes | Monocentridae | *Monocentris* | *Monocentris japonicus* | | 94.4-97.6% | 2-3 |
| | | Trachichthyidae | *Paratrachichthys* | *Paratrachichthys trailli* | sandpaper fish | | |
| | undefined | Opistognathidae | *Opistognathus** | *Opistognathus iyonis** | | 89.9-90.5% | 2-3 |
| | | | | *Opistognathus liturus** | seto-amadai | 89.3-90.5% | 2 |
| | | | | *Opistognathus punctatus** | finespotted jawfish | 81.70% | 5 |
| | | | | *Opistognathus sp.** | | 85.70% | 2 |
| | Uranoscopiformes | Pinguipedidae | *Parapercis* | *Parapercis colias* | New Zealand blue cod | | |
| | | | | *Parapercis decemfasciata* | | 80.90% | 1 |
| | | | | *Parapercis gilliesii* | yellow weaver | | |
| Chondrichthyes | Carcharhiniformes | Carcharhinidae | *Prionace* | *Prionace glauca* | blue shark | | |
| | | Scyliorhinidae | *Cephaloscyllium* | *Cephaloscyllium isabellum* | | | |
| | | Triakidae | *Galeorhinus* | *Galeorhinus galeus* | tope shark | | |
| | | | *Mustelus* | *Mustelus lenticulatus* | spotted estuary smooth-hound | | |
| | | | | *Mustelus manazo* | starspotted smooth-hound | 98.90% | 0 |
| | Hexanchiformes | Hexanchidae | *Notorynchus* | *Notorynchus cepedianus* | broadnose sevengill shark | 98.40% | 0 |
| | Lamniformes | Alopiidae | *Carcharodon* | *Carcharodon carcharias* | great white shark | | |
| | | | *Isurus* | *Isurus oxyrinchus* | shortfin mako shark | | |
| | Squaliformes | Squalidae | *Squalus* | *Squalus acanthias* | spiny dogfish | | |
| | | | | *Squalus suckleyi* | Puget Sound dogfish | 100% | 0 |
